# A unified language model bridging *de novo* and fragment-based 3D molecule design delivers potent CBL-B inhibitors for cancer treatment

**DOI:** 10.1101/2025.11.13.688260

**Authors:** Han Wang, Guanglong Sun, Bowen Zhang, Yang Wang, Bin Xi, Minjian Yang, Chuanyu Liu, Yuyang Ge, Fan Fan, Wei Feng, Yanhao Zhu, Yang Xiao, Yuji Wang, Zhenming Liu, Daohua Jiang, Huting Wang, Wenbiao Zhou, Bo Huang

## Abstract

The rational design of small molecules is central to drug discovery, yet current artificial intelligence (AI) methodologies for generating three-dimensional (3D) molecules are often siloed, focusing on either *de novo* design or fragment-based design. The lack of a holistic framework limits AI’s application across the complex and multi-step pipeline spanning from novel scaffold identification to lead compound optimization, and prevents AI from effectively learning from the entire process. Here, we introduce UniLingo3DMol, a language model for 3D molecular generation, empowered by fragment permutation-capable molecular representation alongside multi-stage and multi-task training strategy. This integrated design enables UniLingo3DMol to seamlessly span both *de novo* and fragment-retained molecular design, demonstrating superior performance over existing generation models in *in silico* evaluations across more than 100 diverse biological targets. We further leveraged UniLingo3DMol in the design of inhibitors targeting CBL-B, a crucial immune E3 ubiquitin ligase and attractive immunotherapy target. This strategy led to a lead compound demonstrating excellent *in vitro* activity and robust *in vivo* anti-tumor efficacy. Our findings establish UniLingo3DMol as a generalized and powerful platform, showing the strong potential to advance AI-driven drug discovery.

## 1. Introduction

The discovery and development of innovative therapeutics remain an exceptionally capital-intensive and time-consuming endeavor, with high attrition rates across multiple complex stages^1^. Artificial intelligence (AI) has emerged as a transformative force, offering unprecedented potential to accelerate key bottlenecks in drug development, including enhanced hit identification^2–8^, efficient virtual screening^9–12^, and optimized lead candidate selection^13–16^.

Drug design methodologies fundamentally diverge into two principal paradigms: ligand-based drug design (LBDD) and structure-based drug design (SBDD)^17^. Each paradigm contributes distinct, yet complementary, knowledge essential for molecular innovation. LBDD operates within chemical space, focusing on chemical validity and the elucidation of structure-activity relationships (SAR), which reveals how systematic structural modifications influence biological activity^18,19^. In contrast, SBDD operates within three-dimensional (3D) space, focusing on ligand-pocket binding efficiency and the delineation of critical non-covalent interactions (NCIs) between ligand and pocket^20^. Drug designers traditionally integrate SAR information from LBDD with binding mode insights from SBDD in an iterative and synergistic fashion^21,22^. They interpret SAR through the lens of ligand-pocket interactions, inferring crucial preferences such as sub-pocket occupancy and the relative importance of specific NCIs. This deepened understanding of binding modes then guides the rational design of new molecules to further enrich SAR and thereby further refine binding modes, a continuous cycle that ultimately yields novel molecule structures with optimal pocket-binding. A classic illustration of this integrated approach is the development of ABT-199 (Venetoclax)^23^, a potent and selective BCL-2 inhibitor, where SBDD-derived atomic structures and LBDD-driven SAR consistently indicated the feasibility and potential benefit of introducing an indole moiety to the ABT-199 scaffold for further sub-pocket occupancy—a feature ultimately confirmed as a key contributor to ABT-199’s potency.

Contemporary AI-assisted drug design (AIDD) efforts have provided support for both LBDD and SBDD tasks independently. LBDD-focused AI molecule generation models, such as CProMG^24^ and MoLeR^25^, effectively capture SAR by leveraging molecular fingerprints or graph representations to predict compound activity and generate novel molecular structures. However, these methods frequently lack explicit consideration of biomolecular structures in 3D space, and often struggle to provide molecules with optimal interactions with binding pockets. Concurrently, SBDD-focused generative models like Pocket2Mol^2^, TargetDiff^4^, Lingo3DMol^8^, and MolCraft^26^ proficiently demonstrate the ability to understand and utilize atomic-level 3D pocket features for generating 3D geometrically complementary ligands. However, despite their effectiveness in capturing plausible binding modes, these models have been reported to produce molecules with flawed two-dimensional (2D) topologies and limited drug-likeness^27^. These challenges underscore a pressing need for AI to achieve a more profound, synergistic fusion of ligand-centric chemical information and 3D binding mode insights, mimicking the sophisticated cognitive processes of human experts. This ambition consequently has catalyzed increased interest in generative AI-driven fragment-based drug design (FBDD)^28^. In this field, researchers often aim to either retain key fragments identified through SAR and innovatively connect them to generate molecules with novel scaffolds, or to grow substituents from a pre-defined scaffold within a binding pocket to optimize interactions. This strategy effectively distills LBDD’s SAR into manageable fragments and their combinations, which are then used as a basis to explore optimal interactions with the pocket in 3D space. Recent advancements, including models like Delete^16^, DeepFrag^29^, and DiffLinker^30^, support single-fragment growth or multi-fragment linking within 3D pockets. However, existing FBDD-focused AI models primarily address fragment-based scenarios and are critically limited in their capability for *de novo* ligand generation within an empty binding pocket. This limitation prevents the crucial synergy where fragment-retained and *de novo* design tasks mutually reinforce each other’s learning, hindering AI’s capacity to truly learn the core human cognition behind integrating SAR and binding mode information.

Bridging this critical gap necessitates a fundamental shift toward a comprehensive AI solution capable of synergistically learning from both pocket-aware *de novo* and fragment-based design. To this end, we introduce UniLingo3DMol, a language model adept at both *de novo* and fragment-based design within a pocket-aware context. We developed a molecular representation which supports fragment permutation, thereby unifying *de novo* generation and fragment-retained generation within the same autoregressive sampling framework. In addition, we employed a multi-stage and multi-task training framework to enhance the model’s learning of chemical plausibility and pocket binding mode validity. These designs engineered our model to internalize crucial binding pocket interactions alongside ligand-centric information, thereby enabling comprehensive molecule generation across both *de novo* and fragment-based scenarios. To support UniLingo3DMol training, we established RComplex database, a protein-ligand complex database created by leveraging high-resolution experimentally determined structures and literature-reported ligands. Through rigorous *in silico* evaluation on 102 targets from Directory of Useful Decoys-Enhanced (DUD-E) dataset^31^, UniLingo3DMol achieved state-of-the-art (SOTA) performance across key metrics of chemical properties, molecular conformations, binding mode quality, and reproducibility of known active compounds in both *de novo* and fragment-based design. Furthermore, demonstrating its practical utility, UniLingo3DMol facilitated the discovery of potent inhibitors for CBL-B, a crucial immune E3 ubiquitin ligase negatively regulating immune cell activity and considered as an attractive immunotherapy target. UniLingo3DMol supports both initial hit compound design and subsequent lead optimization, ultimately yielding a potent CBL-B inhibitor exhibiting nanomolar biochemical activity, potent cellular activity (IC₅₀ 159 nM), and promising *in vivo* efficacy, achieving 76% tumor growth inhibition when combined with a PD-1 antibody in a syngeneic murine model.

## 2. Results

### 2.1. Development of UniLingo3DMol Model

UniLingo3DMol is language model designed for the autoregressive generation of molecules and their conformations directly from an input binding pocket. Its architecture, as illustrated in Figure 1a, comprises a transformer-based encoder-decoder framework. Specifically, UniLingo3DMol employs a 6-layer encoder, a 12-layer decoder, and three ligand heads as its core components. Additionally, a pocket head can be integrated depending on the training stage to perform auxiliary tasks.

**Figure 1.**
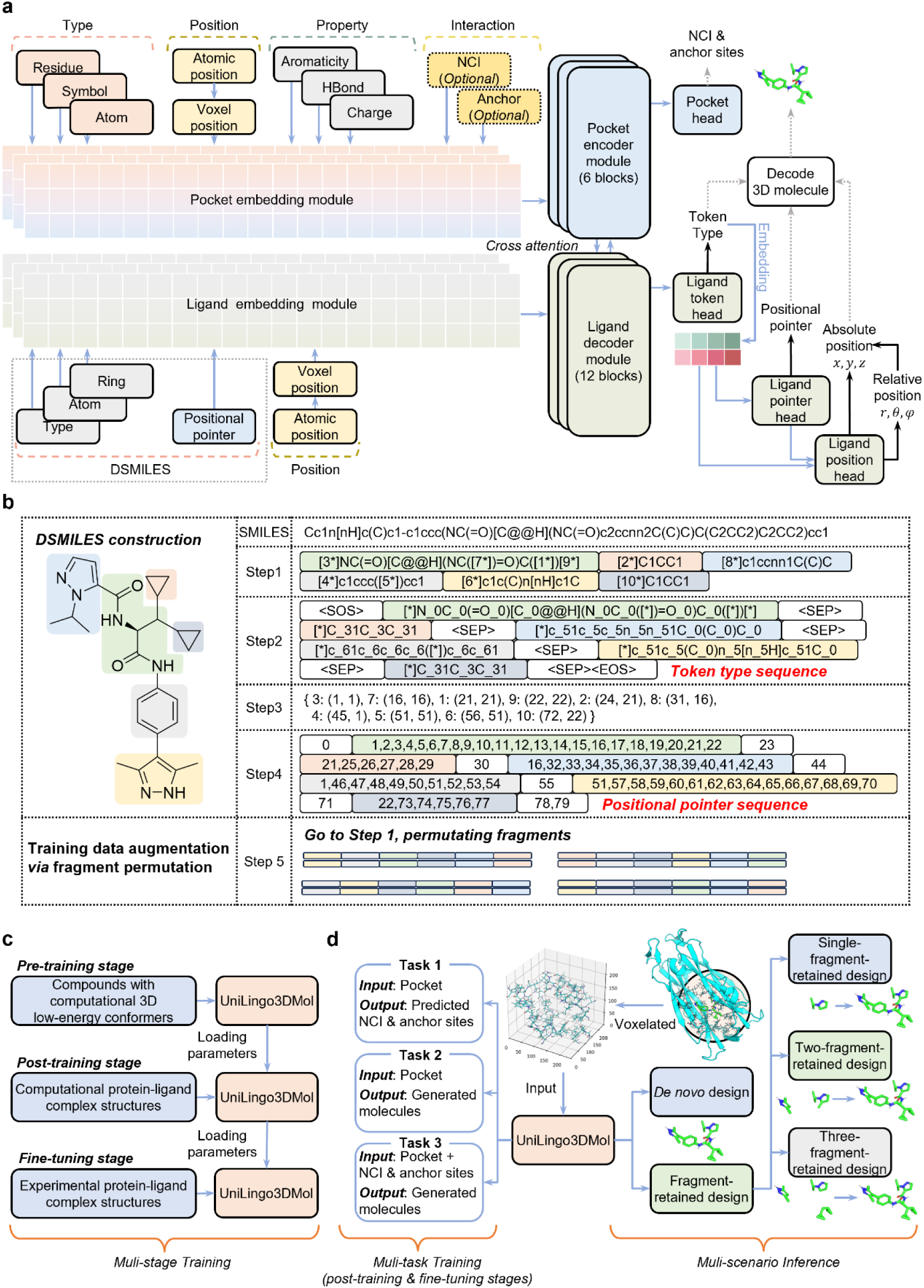
Overview of UniLingo3DMol model development. **(a) UniLingo3DMol architecture.** The model employs a transformer-based encoder-decoder with a 6-layer encoder, 12-layer decoder, three ligand heads and one pocket head. It directly generates molecules and 3D conformations by producing DSMILES token type, positional pointer, and atomic coordinate sequences from an input binding pocket. **(b) DSMILES molecular representation workflow.** This process first involves molecule segmentation into fragments (indicated by colorful blocks). Second, the DSMILES token type sequence is generated, embedding ring information. Third, connection relationships are encoded into a structured dictionary, mapping bond-breaking asterisks with positional indices. Fourth, the positional pointer sequence is constructed. Notably, introducing fragment permutation in Step 2 can create variant sequences of the same molecule, thereby supporting data augmentation during model training. More details are provided in Methods Section 4.2. **(c) Multi-stage training.** UniLingo3DMol undergoes a three-stage sequential training: pre-training (learning basic chemical and conformational rules by using computational low-energy ligand conformers), post-training (learning ligand-pocket binding modes by using force-field accuracy protein-ligand complex structures), and fine-tuning (using experimental accuracy protein-ligand complexes). **(d) Multi-task training and multi-scenario inference.** Three distinct tasks are used in post-training and fine-tuning to thoroughly learn ligand-pocket binding modes. Task 1 is for NCI and anchor prediction; Task 2 is for unconstrained molecular generation; and Task 3 is for conditional generation with NCI and anchor constraints. UniLingo3DMol supports *de novo* design, as well as single-, double-, and triple-fragment-retained molecular design.

#### 2.1.1. Fragment Permutation-Capable Molecular Representation

SMILES syntax^32^, wherein atomic bonding is implicitly derived from token’s neighborhood, presents limitations for autoregressive molecular generation when retaining multiple and unconnected fragments. Specifically, it doesn’t support the representation of these unconnected fragments all placed at the beginning of a sequence and the subsequent autoregressive construction of a larger molecule around them. This limitation consequently hinders the training of both *de novo* generation and fragment-based generation models using a unified framework. To address this gap, we introduce Dual sequence-based SMILES (DSMILES), a unified molecular representation syntax. This syntax (Figure 1b) encodes molecular topology using a dual-information sequence: one specifying token types and the other defining fragment connectivity *via* positional pointers. This explicit representation of connectivity allows DSMILES to accommodate any retained fragments, irrespective of their initial connectivity, by positioning them at the beginning of a sequence.

To fully describe a molecule, beyond its 2D topology encoded by DSMILES, its 3D conformation is represented by atomic coordinates, provided in a separate list associated with each token type.

Based on this unified molecular representation, generating a molecule and its corresponding 3D conformation requires UniLingo3DMol to employ three distinct ligand heads (Figure 1a): a ligand token head for DSMILES token type prediction, a ligand pointer head for positional pointer prediction, and a ligand position head for atomic coordinate prediction. Further details regarding DSMILES are provided in Methods Section 4.2

#### 2.1.2. RComplex Database for UniLingo3DMol Training

Our RComplex database was constructed from a foundation of 48,222 experimentally determined protein–ligand complexes initially retrieved from the Protein Data Bank (PDB)^33^. These experimentally determined structures were subsequently integrated with a vast array of bioactivity data sourced from ChEMBL^34^, PubChem^35^, and GOSTAR^36,37^. This integration process, illustrated in Extended Data Figure 1a and detailed in Methods Section 4.6, constructed active compounds complexed with their binding proteins and classified them into three distinct confidence levels based on their structural relationship to the co-crystallized ligand associated with the PDB ID of their binding protein. Specifically, Level 0 encompasses the co-crystallized ligand itself, offering the highest structural certainty; Level 1 includes compounds with overall structural similarity and substructure overlap; and Level 2 comprises compounds only meeting overall structural similarity. The resulting RComplex database contains 3.7 million protein-ligand complex structures, encompassing a broad spectrum of protein classes (Extended Data Figure 1b), with ∼45% of protein structures (PDB ID) associated with multiple experimentally validated active ligands (Extended Data Figure 1c).

To prevent data leakage, protein targets from the RComplex database and DUD-E were combined and structurally clustered using FoldSeek^38^ with a TM-score cutoff of 0.5 and a sensitivity setting of 7.5. This process generated non-redundant clusters based on tertiary structure homology. Clusters that contained no DUD-E proteins were subsequently designated as the training set, consisting of ∼1.9 million complex structures.

#### 2.1.3. Data Augmentation Empowered Muti-Stage and Multi-Task Training

As illustrated in Figure 1c, UniLingo3DMol underwent complete training through a pipeline that consisted of three sequential stages: pre-training, post-training, and fine-tuning. Each stage employed a unique combination of training data and objectives.

For pre-training, the encoder receives a 3D ligand conformation corrupted by perturbing element types and atomic coordinates (Extended Data Figure 2a). Concurrently, the decoder aims to reconstruct the original 3D conformation. Through this process, UniLingo3DMol acquired foundational knowledge regarding ligand chemical composition and 3D molecular conformation. The training data consisted of an in-house virtual compound library, containing 8 million molecules with conformations generated using ConfGen^19^.

In contrast, the post-training stage shifted the encoder’s input from perturbed ligand conformations to protein binding pockets (Extended Data Figure 2b). This stage leveraged parameters from pre-training and employed Level 1 and 2 data from the RComplex database (Supplementary Section 2.3). A multi-task training strategy was adopted in this stage for robustly learning of the connections between ligand structure and ligand-pocket binding modes. As depicted in Figure 1d, UniLingo3DMol was developed through training on three distinct tasks, each designed to capture ligand-pocket binding mode information from unique perspectives. Task 1 addressed a classification problem focused on predicting NCI and anchor sites within a given pocket structure. NCI sites were defined as residue atoms forming classical interactions (e.g., hydrogen bond donors/acceptors, π-systems) with the co-crystallized ligand in that pocket. Anchor sites comprised pocket heavy atoms within 4 Å of the co-crystallized ligand. Crucially, the co-crystallized ligand was utilized solely for ground-truth label generation during model training, and was not provided as input during Task 1’s prediction phase. Task 2 focused on unconstrained molecular generation within a given pocket, where binding modes were implicitly learned without explicit NCI and anchor site specifications. In contrast, Task 3 enabled conditional molecular generation in a given pocket, directly guided by specified NCI and anchor constraints to facilitate explicit learning of binding modes. Further methodological details are provided in Methods Section 4.3.6.

During the final fine-tuning stage (Extended Data Figure 2b), we preserved parameters obtained after post-training while enhancing them using the Level 0 data in RComplex database. The three-task strategy used in post-training stage was also applied in this stage.

To enable UniLingo3DMol to operate across *de novo* and fragment-based design scenarios, a tailored data augmentation strategy was employed, specifically by including single-fragment-retained and multi-fragment-retained samples in the training dataset. This strategy leverages DSMILES-enabled fragment permutation (Extended Data Figure 2c). Details of augmentation algorithms are provided in Methods Section 4.3.1 and Supplementary Section 2.1.3.

#### 2.1.4. Versatile Input Handling for Diverse Molecular Design Scenarios

Benefiting from its data augmentation and multi-task training strategy, UniLingo3DMol effectively handles diverse input conditions without requiring model adaptation. The model’s encoder processes two types of input: (1) 3D pocket structures without NCI and anchor site annotations, suitable for scenarios with no prior binding knowledge; and (2) 3D pocket structures with explicit NCI and anchor site annotations on residues, ideal when prior active or co-crystallized ligands provide insights into binding modes. Concurrently, UniLingo3DMol’s decoder operates with two distinct input types: (1) an empty input for *de novo* molecular discovery; and (2) user-defined molecular fragments that are preserved during generation, thereby facilitating fragment-based design.

Importantly, the underlying model parameters of UniLingo3DMol are invariant across these diverse applications, with all scenario-specific adaptations handled solely through input modulation.

### 2.2. UniLingo3DMol Model Evaluation

#### 2.2.1. Superior Performance in *De Novo* Molecular Generation

For *de novo* design evaluation, we employed the protein targets from the DUD-E dataset^31^, a widely adopted benchmark in prior studies^2,4,8^ DUD-E comprises 102 diverse protein targets, including kinases, proteases, GPCRs, and ion channels, with each providing, on average, over 200 experimentally validated active ligands. This robust setup enables direct comparison of generated molecules against known active compounds across a broad range of proteins. Thereby, our assessment strategy included a semi-experimental approach to validate the reproduction of known active compounds, as well as the widely adopted computational metrics for molecular 2D topology, 3D conformation quality, and binding modes. Comprehensive metric definitions are detailed in Table 1 and Supplementary Section 2.3.4.

**Table 1.**
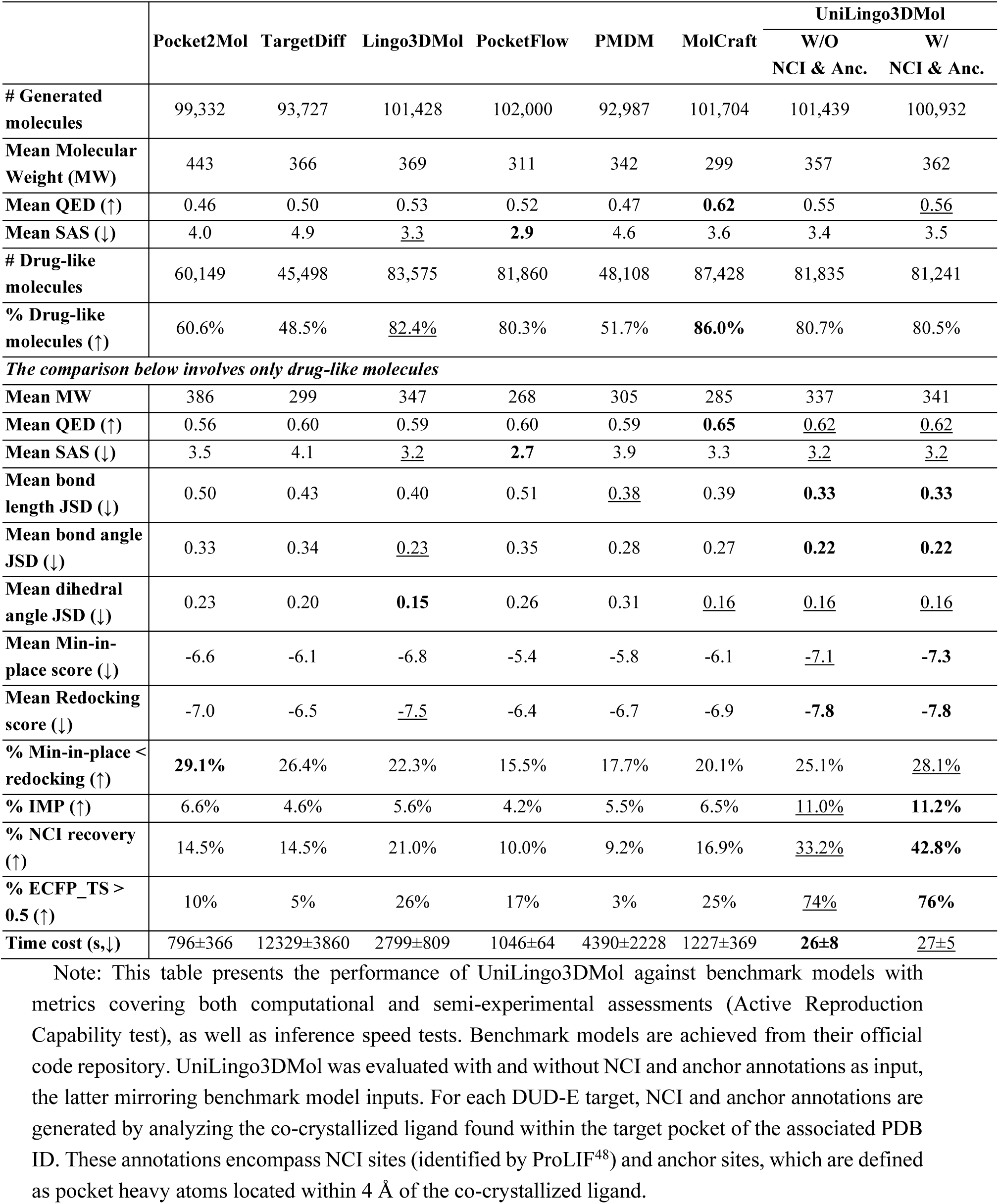
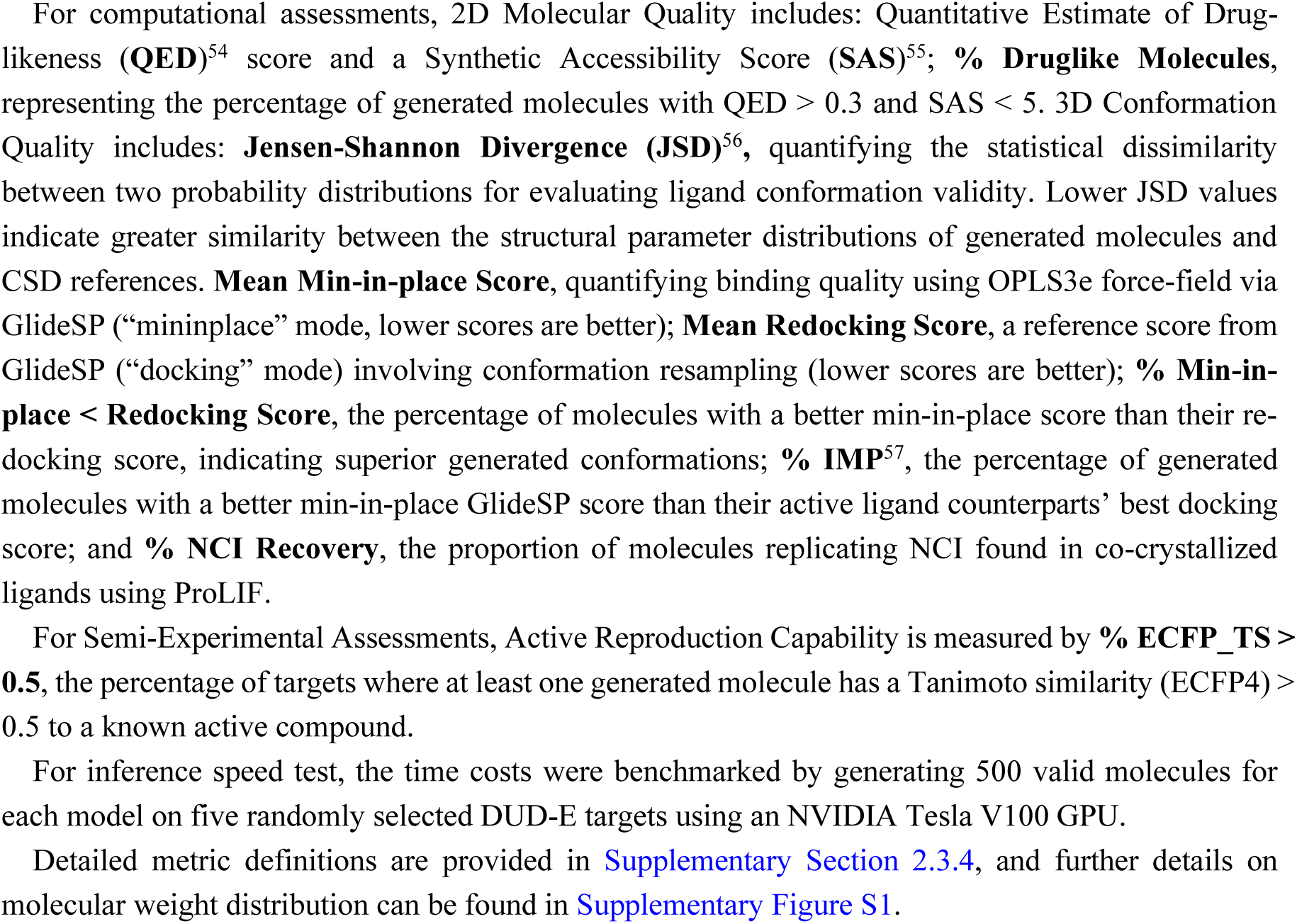
Comparison of generated drug-like molecules on the *de novo* design scenario (N=102). The top two results for each metric are highlighted in bold and <u>underlined</u>, separately.

Six established structure-based generative models, including Pocket2Mol^2^, TargetDiff^4^, Lingo3DMol^8^, PMDM^7^, PocketFlow^5^ and MolCraft^6^, were used as benchmarks. UniLingo3DMol’s performance was assessed under two distinct input conditions: using standard pocket 3D structures without NCI and anchor annotations, mirroring the inputs for benchmark models, and additionally with NCI and anchor annotations. The inclusion of NCI and anchor annotations as input reflects real-world drug design scenarios where NCI and anchor sites, derived from known target protein and its co-crystallized ligand, are often considered crucial for guiding novel binder designs. The NCI and anchor annotations labeled for target proteins in DUD-E are prepared by analyzing the co-crystallized ligand within the target pocket of the associated PDB ID.

For our evaluation, each model generated approximately 1,000 molecules per DUD-E target. We first assessed the 2D quality of these compounds by characterizing their drug likeness. Based on established criteria for drug-like molecules in prior studies^8,39^ (QED > 0.3 and SAS < 5), UniLingo3DMol produced a high percentage (over 80%) of drug-like molecules, comparable to Lingo3DMol and PocketFlow (Table 1). The relatively high drug-likeness observed for MolCraft (around 86%) can be partly attributed to its tendency to produce relatively small molecules, reflected in the lower molecular weights (below 300Da) relative to other models.

Before proceeding to 3D quality assessments, molecules not meeting the drug-likeness criteria were excluded. This filtering step is essential, as molecules with poor drug-likeness can sometimes yield misleadingly favorable scores in force field-based 3D binding evaluations. Consequently, all subsequent analyses focused exclusively on drug-like molecules (QED > 0.3 and SAS < 5). To provide a supplementary view, Extended Data Figure 3 illustrates the distribution of properties of generated molecules within the entire QED-SAS chemical space.

Next, we assessed the 3D conformational quality of the generated drug-like molecules. Ligand-centric conformation validity was evaluated by comparing distributions of bond lengths, bond angles, and dihedral angles against experimentally derived reference data from the Cambridge Structural Database (CSD)^40^. The dissimilarity between these distributions was measured using the Jensen-Shannon Divergence (JSD). UniLingo3DMol demonstrated superior performance compared to most benchmark models, achieving lower JSD values (Table 1). This indicates that our model generates molecular conformations that are more consistent with reference data compared to other models.

Beyond evaluating intrinsic ligand conformation validity, assessing ligand-pocket binding quality is crucial. In this regard, UniLingo3DMol demonstrated superior performance (Table 1). Specifically, our model exhibited superiority in generating conformations with high-quality pocket binding modes, as reflected by Min-in-place Scores. Furthermore, UniLingo3DMol outperformed benchmarks in generating superior binding modes compared to redocked poses (indicated by % Min-in-place < redocking). Additionally, UniLingo3DMol showed enhanced capability relative to benchmarks in generating molecules with better binding modes compared to the co-crystallized ligand (quantified by % IMP). It also surpassed benchmarks in its ability to capture essential pocket-binding NCIs (% NCI recovery), particularly within the scenario without pre-specified NCI and anchor constraints. These collective results robustly underscore UniLingo3DMol’s exceptional ability to generate high-quality 3D molecules with favorable binding modes within the target pocket.

Finally and most importantly, we performed a semi-experimental assessment of active compound reproducibility. This approach allows for a direct comparison of AI-generated molecules against experimentally validated active compounds, thereby addressing the inherent limitations of relying solely on purely computational metrics in AI generative model assessment. Specifically, for each model, we quantified the percentage of targets where at least one generated molecule showed a Tanimoto similarity (ECFP4 fingerprints) greater than 0.5 to any known active compound in the DUD-E dataset (% ECFP_TS > 0.5, Table 1). UniLingo3DMol achieved the highest % ECFP_TS > 0.5, exceeding 70%, represents a substantial advance compared to other models, all of which performed below 35%. As UniLingo3DMol uniquely covers both *de novo* and fragment-based scenarios among all evaluated models, its substantial improvement in *de novo* design implies a notable synergy from its dual-scenario learning.

#### 2.2.2. Enhanced Capability in Fragment-based Molecular Design

In addition to evaluating UniLingo3DMol’s performance in *de novo* design, we employed consistent metrics to assess its capability in fragment-retained design, as detailed in Supplementary Section 1.2. In this scenario, the model needed to generate molecules by incorporating one or more retained fragments. Due to the absence of a readily available evaluation set for this specific scenario, we constructed three dedicated evaluation sets derived from the DUD-E dataset. These sets include: (1) a single-fragment-retained set (144 samples), (2) a two-fragment-retained set (79 samples), and (3) a three-fragment-retained set (19 samples). Details of the method for constructing these evaluation sets are provided in Supplementary Section 2.3.3. In addition, Delete^16^ was used as the baseline method for comparison. Delete is a diffusion-based model for constrained molecule generation in this context, having achieved SOTA performance in fragment-based tasks.

Our evaluation (Supplementary Table S2) demonstrates UniLingo3DMol’s superior performance over the benchmark method in both computational and semi-experimental assessments across all three evaluation sets, including single-, two-, and three-fragment retained scenarios. UniLingo3DMol doubled the percentage of drug-like molecules compared to Delete. It also achieved lower JSD scores across the majority of conformational geometry metrics (bond length, bond angle, and dihedral angle). For binding poses, UniLingo3DMol consistently showed lower min-in-place scores. Additionally, it produced a substantially higher percentage of targets with ECFP_TS > 0.5, signifying improved active compound reproducibility.

### 2.3. UniLingo3DMol-supported Discovery and Optimization of CBL-B Inhibitors

To thoroughly demonstrate the practical application of UniLingo3DMol in drug design, we systematically elaborated its pivotal role in the effective design of CBL-B inhibitors. The overall workflow is illustrated in Figure 2. Generally, we began with a crystal structure (PDB: 8GCY), which contains CBL-B complexed with a known active compound (Figure 2a). Given that UniLingo3DMol demonstrated overall superiority when utilizing pockets with NCI and anchor site labels in our *in silico* evaluation, we extracted such information from the crystal structure and used it as input for the molecule generative model (Figure 2b). The entire process was comprised of a two-fragment-based generation targeting novel core scaffold discovery (Figure 2c), and a single-fragment-retained generation for compound optimization (Figure 2d). Further details on each phase are provided in the subsequent subsections.

**Figure 2.**
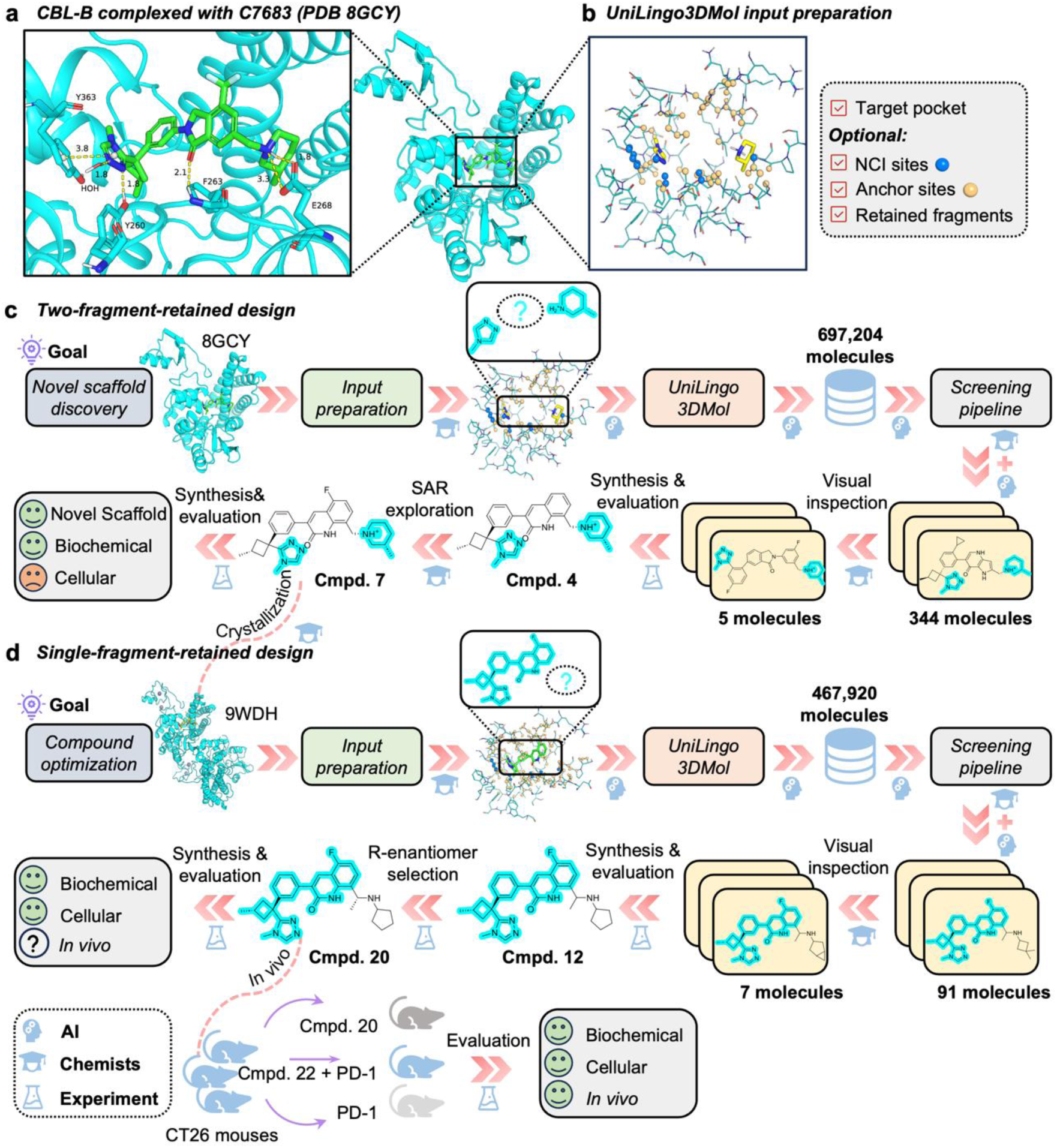
Schematic diagram of UniLingo3DMol-supported CBL-B inhibitor discovery and optimization. **(a)** Structure of CBL-B active binding pocket complexed with C7683. **(b)** Checklist of inputs for UniLingo3DMol. **(c)** Workflow for novel scaffold design. **(d)** Workflow for lead compound optimization.

#### 2.3.1. Novel Scaffold Discovery

The previous study^41^ reported a series of triazole compounds inhibiting the E3 ligase activity of CBL-B for cancer immunotherapy. Building upon this foundational research, we employed UniLingo3DMol model for the design of novel small-molecule CBL-B inhibitors. Specifically, we utilized the crystal structure of CBL-B complexed with an inhibitor C7683 (PDB: 8GCY) as the start point for our molecule generation (Figure 3a). To facilitate a clearer structural description and enable subsequent molecular comparisons, C7683 was divided into four distinct regions: A, B, C, and D (Figure 3a). The methyl triazole moiety in Region A was deemed critical for the inhibition activity in the reported study^41^ and therefore was retained during our molecule generation process. In addition, given its exposure to the solvent-exposed region, the methyl piperidine group in Region C was also conserved in this initial design phase to streamline the chemical space exploration. The amino acid residues of CBL-B within a 6 Å of C7683 were used as the pocket. The pocket structure and the two conserved fragments (Regions A and C) were inputted into UniLingo3DMol for molecule generation.

**Figure 3.**
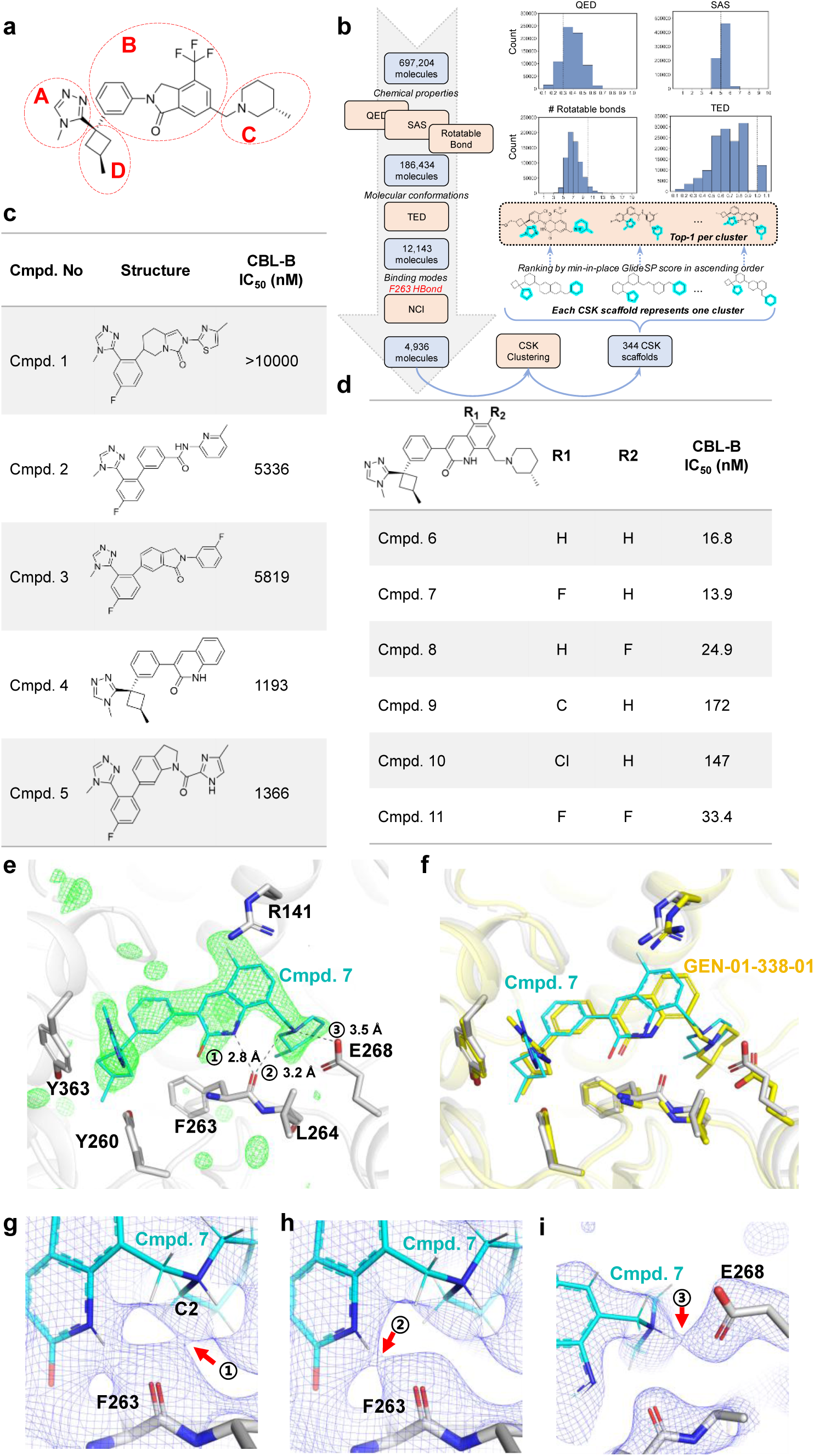
Design strategy and initial compound generation. **(a)** Schematic representation of C7683 (from PDB: 8GCY) divided into four distinct regions (A, B, C, and D). Regions A (methyl triazole) and C (methyl piperidine) were conserved during the initial generative process. **(b)** Schematic diagram illustrating the integrated AI-driven molecular generation and screening pipeline employed for the discovery of novel hit compound scaffolds. In this initial discovery phase, target protein pocket structures served as the input for AI-guided molecule generation. The multi-step screening process involved: (1) drug-likeness triage (QED, SAS, number of rotatable bond); (2) torsion energy screening (TED^39^); (3) binding mode selection (hydrogen bond with F263); (4) diversity (CSK^42^ clustering); and (5) overall binding quality (GlideSP min-in-place scoring). **(c)** Biochemical activity results for the initial five synthesized compounds (A, B, D regions only). **(d)** Biochemical activity and SAR data for Cmpd. 6 and its derivatives (R1/R2 substitutions). **(e-i)** Co-crystal Structure of Cmpd. 7 with CBL-B and NCI Analysis. **(e)** X-ray crystallographic conformation of Cmpd. 7 binding within the CBL-B active site. The F_obs_-F_calc_ omit map, contoured at 3.0 σ (green mesh), confirms the presence of Cmpd. 7. This unbiased omit map was generated by excluding Cmpd. 7 from the crystallographic model (occupancy set to zero) followed by refinement. Cmpd. 7 is shown in cyan stick representation within the binding pocket of CBL-B (white cartoon). Three key NCIs are numerically labeled with corresponding interatomic distances indicated. **(f)** Superposition of the experimentally determined conformation of Cmpd. 7 (cyan) and its UniLingo3DMol model-generated counterpart GEN-01-338-01 (yellow). Our X-ray crystallography determined Cmpd. 7-CBL-B complex is shown with the protein in white and Cmpd. 7 in cyan. The CBL-B structure from PDB 8GCY (yellow) is superimposed with our complex. **(g)** The non-classical hydrogen bond between the carbonyl oxygen of F263 backbone and the C2 atom of the 3-methylpiperidine moiety of Cmpd. 7. **(h)** The conventional hydrogen bond between the carbonyl oxygen of F263 backbone and the nitrogen atom on the quinoline moiety of Cmpd. 7. **(i)** The salt bridge between the protonated nitrogen atom of the 3-methylpiperidine of Cmpd. 7 and the side chain of E268. The 2F_o_-F_c_ electron density maps are shown around the interaction site at contour levels of 0 σ **(g)**, 0.35 σ **(h)**, and 1.3 σ **(i)**, respectively. The bond critical points (BCPs) indicating NCIs are labeled with red arrows.

The UniLingo3DMol model generated a set of ∼700k molecules which were subsequently subjected to a multi-stage screening workflow (Figure 3b; Supplementary Section 1.4.1). This screening workflow encompassed several stages: (1) Preliminary drug-likeness filtering was performed using QED, SAS, and rotatable bond counts. (2) Conformational screening utilized TED^39^ to filter low-strain conformations. (3) Binding mode selection focused on recovering the key hydrogen bond between reference C7683 and F263 of the pocket. (4) CSK^42^ scaffold-based clustering was employed to ensure structural diversity. (5) Finally, from each CSK-based cluster, the top molecule was selected based on its GlideSP score (min-in-place). The top 344 candidates, as detailed in the Supplementary Materials, underwent rigorous visual inspection which focused on assessing their interactions within the binding pocket, the novelty of their molecular scaffolds, and their synthetic feasibility. As shown in Extended Data Figure 4, we identified several designs reflecting classical motifs found in previous patents, such as trifluoromethyl-isoindolin and pyrrolopyridine derivatives in Region B; we also identified various novel moieties within this region. After excluding potentially toxic structures (e.g., three-membered fused ring systems or certain ketones) and those presenting synthetic challenges, five molecules were selected for chemical synthesis and subsequent biochemical activity testing (Extended Data Figure 4). To minimize synthetic complexity and cost, only the A, B, and D regions of these selected compounds were synthesized. As presented in Figure 3c, Cmpd. 2, Cmpd. 3, Cmpd. 4 and Cmpd. 5 demonstrated micromolar biochemical activity. Of these, Cmpd. 4 demonstrated the strongest potency and thus was chosen for the reintroduction of the C region, yielding Cmpd. 6 (Figure 3d). This compound demonstrated high biochemical potency, exhibiting an IC_50_ of 16.8 nM. To further enhance the biochemical activity of Cmpd. 6, a preliminary SAR study was conducted, focusing on the substitution of the R1 and R2 groups of the 2-quinolone moiety in Region B. As evident from Figure 3d, these simple substitutions did not lead to a substantial improvement in biochemical activity. However, the introduction of a fluorine atom at the R1 position (Cmpd. 7) resulted in a modest enhancement of biochemical activity. Despite the promising biochemical activity, Cmpd. 7 exhibited suboptimal cellular activity: it shows low activity of stimulating Jurkat cell for IL-2 release (3.5 folds change relative to anti CD3/28 at a 370 nM of Cmpd. 7).

#### 2.3.2. Lead Compound Optimization

To conduct optimization on the basis of Cmpd. 7, we put our subsequent efforts towards a systematic investigation of Region C, given that Region A was considered structurally essential and comprehensive explorations of Regions B and D had already been conducted. To facilitate the optimization of Region C, the co-crystal structure of Cmpd. 7 in complex with CBL-B was determined at a resolution of 2.8 Å (Figure 3e; Extended Data Table 1). This experimental structure demonstrated high consistency with the computationally generated conformation of the counterpart molecule (Figure 3f), thereby validating the 3D conformational generative capabilities of our UniLingo3DMol model. Analysis of the bond critical points (BCPs) within the experimental electron density map of the Cmpd. 7-CBL-B complex revealed the presence of a non-classical hydrogen bond between the carbonyl oxygen of the F263 backbone and the C2 atom of the 3-methylpiperidine moiety of Cmpd. 7 (Figure 3g). Furthermore, the carbonyl oxygen of the F263 backbone was also involved in a conventional hydrogen bond with a nitrogen atom on the quinoline moiety of Cmpd. 7 (Figure 3h). Concurrently, the 3-methylpiperidine of Cmpd. 7 established a salt bridge with E268 side chain *via* its protonated nitrogen atom (Figure 3i). These observations of the NCIs provided guidance for the optimization of the Region C: specifically, to replace the identified non-classical hydrogen bond with a classical hydrogen bond while preserving other key NCIs.

For the optimization of Region C, the UniLingo3DMol model was again employed. For this round of generation, our Cmpd. 7-CBL-B complex structure was used as basis with amino acid residues within a 6 Å of Cmpd. 7 defined as the binding pocket and the A, B, and D regions of Cmpd. 7 retained. This generative process yielded ∼500k molecules, which were subsequently screened based on a screening pipeline (Figure 4a) adapted from the preceding generation round. Specific modifications to this screening pipeline included the usage of the mandatory fulfillment of maintaining two conventional hydrogen bonds with F263 and a salt bridge with E268, which are observed in Cmpd. 7-CBL-B crystal structure.

**Figure 4.**
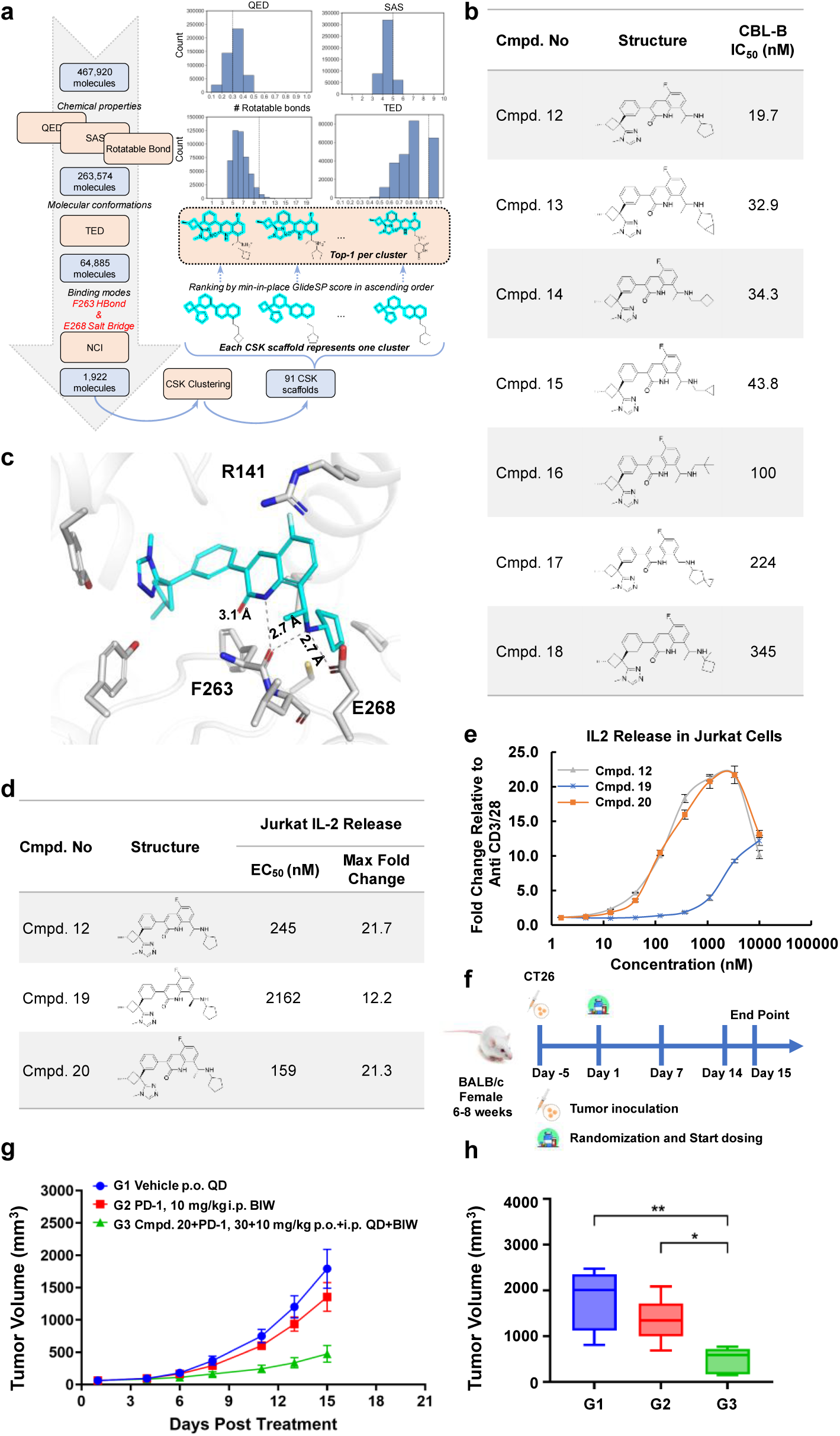
AI generated conformation, cellular activity and *in vivo* efficacy of Cmpd. 20. **(a)** Schematic diagram of the AI-powered molecule generation and screening pipeline utilized for lead compound optimization. This pipeline is adapted from preceding rounds (Figure 3b), with key criteria included in binding mode selection to maintain two hydrogen bonds with F263 and a salt bridge with E268. **(b)** Biochemical activity results for the seven selected compounds from the C region optimization round. **(c)** UniLingo3DMol model-generated 3D conformation of Cmpd. 20 (R-enantiomer of Cmpd. 12) bound within the CBL-B active pocket. Key NCIs, including hydrogen bonds and the salt bridge involving F263 and E268, are indicated with dashed lines. **(d)** Cellular activity and chiral separation results for Cmpd. 20 (R-enantiomer of Cmpd. 12). **(e)** Dose-Response for IL-2 Secretion in Jurkat Cells. Data are presented as mean ± SD from three independent replicates. **(f)** Schematic diagram of the *in vivo* study of Cmpd. 20 in CT26 tumor-bearing mice. **(g)** *In Vivo* Anti-Tumor Efficacy in a CT26 Mouse Model. CT26 tumor-bearing mice were treated *via* oral administration with vehicle (daily), Cmpd. 20 (daily at 30 mg/kg), anti-mouse PD-1 (anti-mPD-1) antibody (bi-weekly), or their combination. Values represent mean tumor volume ±SEM (n=5) mice per group. **(h)** Comparison of tumor volume at study endpoint. Tumor volumes on Day 15 of the study were compared. Comparison made between vehicle-treated group (G1) and Cmpd. 20 plus anti-mPD-1 (G3) is to indicate the efficacy of Cmpd. 20 in a combination treatment; comparison made between anti-mPD-1 (G2) and G3 is to indicate the enhanced efficacy due to the addition of Cmpd. 20 to anti-mPD-1 treatment. A two-sided T-test was employed for statistical analysis, with significance levels indicated as: *p ≤ 0.05, **p ≤ 0.01 (n=5 mice per group).

From the top 91 filtered compounds (Supplementary Materials), seven were selected for synthesis (Figure 4b) *via* visual inspection. This visual selection prioritized compounds satisfying the NCI requirements, while also ensuring the introduction of diverse hydrophobic groups to engage the L148, C289, and P72 regions. For NCI satisfaction, it was observed that a protonated basic amine could simultaneously fulfill the requirements about forming NCIs with E268 and F263 (Figure 4c).

Although the UniLingo3DMol-generated 3D molecular conformations described specific chirality, the selected compounds were initially evaluated for biochemical activity as racemic mixtures due to the resource-intensive nature of chiral separation. The most active compound, Cmpd. 12, was subsequently subjected to chiral separation and cellular activity tests (Figure 4d-4e). The results confirmed that the R-enantiomer (Cmpd. 20), consistent with the UniLingo3DMol-predicted conformation (Figure 4c), exhibited superior cellular activity with an IC_50_ of 159 nM.

Further *in vivo* efficacy study of Cmpd. 20 was undertaken (Figure 4f). In a murine CT26 syngeneic tumor model, oral administration of Cmpd. 20 at 30 mg/kg once daily, in combination with a PD-1 antibody (bi-weekly), resulted in a robust anti-tumor efficacy (Figure 4g-4h) with a tumor growth inhibition (TGI) of 76%. Safety assessments (Extended Data Table 2) indicated a low risk for CYP inhibition/induction and hERG inhibition. Furthermore, Cmpd. 20 exhibited good animal tolerability, with no observed adverse reactions. These collective findings strongly support the designation of Cmpd. 20 as a promising lead compound for further therapeutic development.

## 3. Discussion

In this study, we introduced UniLingo3DMol, a unified deep learning framework for molecular generation that seamlessly integrates 3D pocket-aware *de novo* design and fragment-based approaches. This end-to-end model features a flexible DSMILES molecular representation, a three-stage multi-task training pipeline, and a transformer-based architecture adaptable to diverse drug discovery scenarios. Benchmarking against SOTA models demonstrates UniLingo3DMol’s superior performance in generating drug-like molecules with robust binding modes and high reproducibility. Its practical utility was validated by the discovery and optimization of a potent CBL-B inhibitor, exhibiting nanomolar potency and robust *in vivo* efficacy in a murine tumor model.

In this discussion section, we first analyzed key factors in our training strategy contributing to UniLingo3DMol’s performance, and then discussed our thinking on the best practice of applying generative AI in drug design by summarizing our practice in UniLingo3DMol in CBL-B inhibitor discovery. At last, we discussed the limitations of UniLingo3DMol associated with future research directions considering dynamic data from multi-perspectives.

To elaborate on the key factors contributing to UniLingo3DMol’s performance, we conducted a series of ablation analyses as below.

UniLingo3DMol employs a three-stage training pipeline designed to overcome the data imbalance between chemical space and 3D binding modes. Pre-training on a large virtual compound library establishes fundamental chemical rules and diversity. A subsequent post-training stage uses a medium-sized dataset of computationally derived active ligand-protein complexes to teach molecular interaction principles. Finally, fine-tuning on a smaller set of high-quality experimental complexes aligns the model’s output with real-world binding environments, with post-training serving as a bridge to practical ligand-binding modes. Our ablation studies (Supplementary Table S3) underscore the importance of each stage: while the absence of diverse chemical space learning (i.e., without pre-training) did not harm the chemical validity of generated molecules, it directly led to a drop in known active compound reproducibility (% ECFP_TS>0.5) and 3D pocket-binding quality (Min-in-place GlideSP score, % Min-in-place < redocking, % IMP, % NCI recovery). This implies that the model had a limited chemical space for constructing valid molecules and exploring optimal binding modes within the pocket. Omitting the post-training stage resulted in an abrupt and illusory transition from chemical space to ligand-pocket binding, where apparent good 3D pocket-binding quality was decoupled from poor reproducibility of known active compound.

Our multi-task training strategy is designed for robust ligand-pocket binding learning, crucial due to limited data and its focus across all three tasks on recognizing binding modes. Task 1 identifies specific pocket-ligand interaction regions. Task 2 implicitly learns binding by generating molecules without explicit NCI and anchor site guidance, while Task 3 explicitly learns with NCI and anchor site information. Ablating Task 1 (NCI and anchor site recognition) severely impacted both 3D pocket-binding quality and known active compound reproducibility (Supplementary Table S4). This integrated understanding was further underscored by superior end-to-end inference. Specifically, compared to the end-to-end generation without explicit NCI guidance (Task 2), cascaded inference (Task 1 followed by Task 3) produced deceptively good 3D pocket binding but poor known active reproducibility (Supplementary Table S5).

Benefiting from the DSMILES syntax, which supports free fragment permutation and thereby allows multiple retained fragments to be reordered to the sequence’s beginning, our method can train *de novo* and fragment-based generation using a unified training framework that incorporates both sample types. Our ablation study (Supplementary Table S7) showed that when fragment-retained data was excluded from training, both 3D pocket-binding quality and known active compound reproducibility for *de novo* design dropped notably. This trend indicates that the *de novo* generation ability benefited from the addition of fragment-based training data, suggesting that UniLingo3DMol achieved a synergistic fusion of ligand-derived fragment insights and 3D pocket binding mode information.

In addition, UniLingo3DMol model’s understanding on the intricate correlation between ligand structures and their respective protein pocket binding modes can be visualized through its cross-attention mechanism. For instance, applying our trained decoder layer to process CBL-B with its co-crystallized ligand (PDB ID: 8GCY, Extended Data Figure 5a) enables us to delineate specific regions of the binding pocket that elicit significant “attention” from the ligand. As illustrated in Extended Data Figure 5b, the residues garnering substantial attention from the entire ligand include those involved in polar NCIs (e.g., E268 forming a salt bridge, and F263 forming a hydrogen bond), and those instrumental in defining the pocket’s overall shape and engaging in nonpolar NCIs (e.g., M366, A262, and L264). Moving beyond a holistic view, an examination from the perspective of ligand fragments demonstrates a varied attention pattern for distinct fragments (Extended Data Figure 5c-5f), where these patterns highlight residues directly surrounding the fragment and those engaged in classical NCIs with the fragment. These correlations collectively demonstrate UniLingo3DMol’s understanding of ligand structure and its interactions within the protein pocket.

Our CBL-B inhibitor discovery campaign highlights that generative AI models, despite their power, function as sophisticated tools that augment—rather than replace—human expertise in drug design. Their success critically hinges on medicinal chemists providing strategic guidance, particularly through the definition of retained fragments. For instance, the retention of a methyl triazole in Region A and a methyl piperidine in Region C during novel scaffold generation was instrumental in identifying Cmpd. 4 and 7. To quantify the impact of such guidance, we systematically varied the retained fragments (Extended Data Figure 6a). We observed that exploring a larger chemical space by retaining only Region A or C significantly reduced the probability of generating the desired scaffold. This trend was further corroborated in our search for novel structures within Region A. Specifically, by retaining Region C and either full or partial Region B (Extended Data Figure 6b), our model successfully generated a 2-piperidinone moiety to replace the methyl triazole in Region A, a design consistent with recently reported active compounds^43^. Conversely, exploring a broader chemical space with only Region C retained struggled to produce this desired design within reasonable sampling efforts. Therefore, effective utilization of UniLingo3DMol requires drug designers to strategically vary retained fragments to achieve optimal design outcomes.

A crucial future direction for UniLingo3DMol’s enhancement lies in its adaptability to dynamic discovery environments. This is particularly vital in real-world drug design projects where knowledge continuously accrues; for instance, as SAR insights evolve for a scaffold, optimal binding fragments for specific sub-pockets may be updated. Therefore, establishing a dynamic, real-time, and retraining-free mechanism to seamlessly integrate these updated fragments into molecule generation is paramount. To this end, the integration with retrieval-augmented generation (RAG) presents a promising direction. To demonstrate the feasibility of this approach, we constructed a straightforward prototype named Molecular Retrieval-Augmented Generation (MRAG). This prototype retrieves relevant molecular fragments from an external fragment database, guided by key protein residues. These retrieved fragments are then positioned within an appropriate sub-pocket and explicitly considered as retained fragments during the subsequent molecule generation. While MRAG notably improved the 3D pocket-binding quality of generated molecules, a limitation observed was a reduction in diversity. This could be attributed to the relatively small size of the fragment library supporting our MRAG prototype, which was designed solely for proof-of-concept purposes. Further details about MRAG implementation and evaluations are provided in Supplementary Section 2.2.

Beyond dynamic fragment updates, other advanced methods necessitating model retraining, such as reinforcement learning (RL), represent a powerful direction for incorporating broader SAR information encompassing not only fragment structures but also diverse activities and even pharmacokinetic properties. By designing reward functions that integrate these diverse objectives (potentially leveraging predictions from dedicated property models), the RL framework can iteratively refine UniLingo3DMol, guiding it toward more optimal regions of the multidimensional property space. This approach is expected to provide a more powerful and direct means for multi-parameter lead optimization compared to simple filtering after generation.

Another critical perspective for enhancing UniLingo3DMol’s dynamic environment perception involves accounting for protein flexibility. A key limitation of current structure-based generation methods, including UniLingo3DMol, is their reliance on static protein structures. These methods often overlook that proteins are dynamic molecules undergoing conformational changes. To explicitly integrate protein flexibility and enhance the binding adaptability of generated molecules, a promising strategy involves leveraging advanced protein conformation sampling tools (such as BioEmu^44^) to construct an ensemble of protein conformations. While UniLingo3DMol inherently takes static input structures, its high inference speed allows for using multiple representative conformations from this ensemble as distinct inputs for molecule generation, thereby exploring flexible binding sites.

In summary, this work represents a major step towards a highly unified AI solution capable of effectively learning and applying the combined wisdom of pocket-aware *de novo* design and fragment-based design, thereby paving the way for more rational, efficient, and ultimately successful drug design.

## 4. Methods

We presented UniLingo3DMol, a unified language model for structure-based drug design applicable to multiple drug discovery scenarios, including *de novo* design and fragment-retained design. Two key innovations are introduced in UniLingo3DMol: (1) DSMILES, a novel molecular notation syntax that enables unified representation across diverse molecular generation scenarios. (2) A data augmentation empowered multi-stage and multi-task training strategy that enhances model adaptability without requiring task-specific architectural adjustments. The comprehensive architecture of UniLingo3DMol was detailed in Figure 1.

### 4.1. Problem Definition

UniLingo3DMol tackles 3D molecular generation by probabilistically modeling pocket–ligand interactions. For *de novo* design, the objective of the model is to optimize parameters *θ* that satisfy the probability distribution Mol ∼ *P_θ_*(Mol ∣ Pkt; *θ*). For fragment-retained design, this formulation is extended to Mol ∼ *P*_*θ*_(Mol ∣ Pkt, *L*_partial_; *θ*). Notably, the model maintained parameter consistency across scenarios through a data augmentation empowered unified multi-stage and multi-task learning strategy.

In this study, a binding pocket is defined as residues within 6 Å of the ligand in protein-ligand complexes. Formally, Pkt = (*p*_1_, *p*_2_, …, *p*_*i*_), where each *p_i_* represents a heavy atom characterized by nine physicochemical descriptors: *p*_*i*_ = 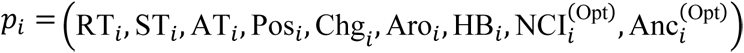. These descriptors are systematically defined as follows: RT (residue type), ST (chemical symbol type), AT (residue-specific atom name), Pos (3D spatial coordinates), Chg (atomic formal charge), Aro (atomic aromaticity), HB (hydrogen bond capability), NCI (non-covalent interaction site indicator), and Anc (anchor site marker indicating proximity ≤ 4 Å to ligand atoms). The “(Opt)” superscript denotes optional features depending on data availability.

The ligand molecule is encoded as Mol = (*m*_1_, *m*_2_, …, *m*_*j*_), where each token *m_j_* contains five distinct components, formulated as *m*_*j*_ = (DT_*j*_, ST_*j*_, CRT_*j*_, Pos_*j*_, Ptr_*j*_): DT (DSMILES type) categorizing the type of the DSMILES token, ST (chemical symbol type) identifying its chemical symbol, CRT (chemical ring system classification) tracking its chemical ring system classification, Pos (3D spatial coordinates) encoding its 3D spatial coordinates (*x*, *y*, *z*), and Ptr (pointer to other tokens) defining connectivity relationships by referencing indices of other tokens linked to *m_j_*. Partial structural constraints *L*_partial_ = (*l*_1_, *l*_2_, …, *l*_*k*_) adopt this identical five-component schema, maintaining representational consistency with full molecular structures while enabling compatibility through shared feature space alignment, thereby ensuring seamless integration of fragmentary chemical constraints into the molecular generation process.

### 4.2. DSMILES: A Unified Molecular Representation Syntax for Drug Discovery

We proposed DSMILES, a molecular representation syntax that enables unified cross-scenario compatibility for UniLingo3DMol. This syntax decomposes molecular structures into chemically meaningful fragments represented in standard SMILES^32^, while explicitly encoding inter-fragment connectivity. The resulting representation preserves molecular topology in sequence format, effectively bridging the advantages of language models and graph neural networks.

DSMILES syntax is fundamentally fragment-based. Its molecular fragmentation employs chemically informed rules to delineate a molecule into its constituent ring systems, sidechains, and linkers. A ring system is defined as any interconnected assembly of single, fused, or spiro-linked rings. To ensure the inclusion of meaningful functional groups, any ring-connected moieties possessing fewer than five heavy atoms are assimilated into its connected ring system. Structural moieties connecting to only one ring system are classified as sidechains. Conversely, structural moieties connecting to more than one ring system are defined as linkers.

DSMILES is comprised of a token type sequence and a positional pointer sequence. Beyond standard SMILES^32^, which encodes element type, aromaticity, bond order, and chirality, DSMILES extends its definition in two perspectives (Figure 1b): (1) Atomic environment descriptors encoding localized structural contexts (e.g., “C_3” for aliphatic ring carbons, and “c_6+6” for fused aromatic systems). (2) Explicit connection encoding relationship between different fragments. The former corresponds to the token type sequence, while the latter corresponds to the positional pointer sequence. These two features enabled comprehensive support for various scenarios through simultaneous preservation of fragments and their topological relationships. Extended Data Table 3 provides complete documentation of valid DSMILES token types.

Figure 1b illustrates the workflow of generating the above dual sequences using DSMILES. Firstly, the molecule is segmented into fragments based on predefined chemical rules, while connectivity between fragments is maintained. Connections between fragments follow a specific convention: a bond-breaking site (denoted by “*”) marked with an odd number *n* must be connected to the site marked with the subsequent even number *n* + 1. For instance, the marker “[1*]” is specifically connected to “[2*]”. Secondly, the DSMILES token type sequence is generated, embedding ring information in each token. Thirdly, connection relationships are encoded into a structured dictionary, where each bond-breaking site is mapped to an ordered tuple of positional indices. This tuple includes the index of the current asterisk in the DSMILES token type sequence and the index of its connected asterisk in the sequence. To handle temporal dependencies in sequential representations, if an asterisk points to an index that has not yet appeared in the pointer sequence, it temporarily points to itself. Finally, the positional pointer sequence is constructed, where asterisk values are derived from the dictionary and non-asterisk tokens refer to their respective sequence indices. Notably, DSMILES allows data augmentation through fragment order permutation in the initial step, enabling multiple valid sequence representations for identical molecular structures via variable fragment ordering.

### 4.3. Data Augmentation Empowered Multi-Stage and Multi-Task Training Strategy

#### 4.3.1. Data Augmentation

In practical drug development, three distinct molecular design scenarios are identified: (1) *de novo* design, (2) single-fragment-retained design, and (3) multiple-fragment-retained design. This diversity presents significant challenges for developing unified computational models capable of addressing all three scenarios. The adoption of DSMILES syntax enables UniLingo3DMol to successfully address 3D molecular design requirements across these diverse scenarios. Specifically, to satisfy the requirement of UniLingo3DMol for multi-scenario 3D molecular generation, it is necessary to implement scenario-specific training data augmentation algorithms.

From a data engineering perspective, these scenarios are categorized into two principal groups: (1) *de novo* design and single-fragment-retained design, and (2) multiple-fragment-retained design. The first group shows consistent connectivity patterns, with each subsequent fragment linking to prior ones in a strict sequential order, though not necessarily to an adjacent fragment. This regularity enables the use of a universal augmentation algorithm (Supplementary Algorithm S3), which preserves pattern coherence while allowing stochastic fragment assembly.

In contrast, the second group exhibits more complex connectivity mechanisms. Here, later fragments in the sequence might remain unconnected to preceding fragments until subsequent connection opportunities occur. The number of candidate connections to be determined varies from two to four to meet different drug development requirements. These distinct connectivity patterns necessitate the use of a specialized augmentation algorithm (Supplementary Algorithm S4). Further details are described in Supplementary Section 2.1.3.

#### 4.3.2. Pre-training Stage

The complete training pipelines consists of three sequential stages: pre-training, post-training, and fine-tuning stages, each with distinct data requirements. For the pre-training stage, we used an in-house virtual compound library containing over 20 million commercially accessible structures, which are typically employed in virtual screening and cover a broad chemical space of ring systems, functional groups, and molecular weights. Low-energy conformers are generated for each molecule using ConfGen^45^. Following a procedure similar to Lingo3DMol^46^, we also applied a filtering process that removed complex rings and retained molecules with fewer than three consecutive flexible bonds, yielding a refined set of 12 million molecules.

During the pre-training stage, each molecule was converted into its DSMILES representation and fed into the decoder, enabling UniLingo3DMol to learn fragment composition and connection rules at scale. We also perturbed 3D molecules including atomic types and spatial coordinates, and then fed the perturbed molecule into the encoder. To obtain more training data, Supplementary Algorithm S3 was used for data augmentation. UniLingo3DMol employed teacher-forced autoregressive reconstruction to optimize dual recovery objectives: accurate restoration of perturbed molecular structures to their original DSMILES representations while preserving both topological connectivity and conformational geometry.

#### 4.3.3. Post-training Stage

As described in the pre-training stage, the data fed into the encoder was perturbed molecules rather than pockets, which deviated from the practical usage scenario. To align this bias, we introduced a post-training stage. At this stage, UniLingo3DMol loaded the parameters obtained from the pre-training stage. Due to the scarcity of protein-ligand complex data with experimental accuracy, we calculated a large number of complex data with force-field accuracy through Glide^47^ to increase the data scale (Supplementary Section 2.3.1). To obtain binding pockets to generate pocket-ligand complexes for aligning UniLingo3DMol, we extracted residues around the ligand within the range of 6 Å for each protein-ligand complex. More importantly, to prevent data leakage during training, a clustering-based filtering approach was implemented. Specifically, the 102 protein targets from the DUD-E evaluation database were combined with the protein structures from our dataset. This combined pool was then clustered using FoldSeek^38^, with a TM-score cutoff of 0.5 to define a cluster. All collected protein structures that clustered with any of the DUD-E targets were subsequently removed from our final dataset, resulting in the exclusion of 10,224 proteins. Furthermore, the NCI analysis was performed for each pocket-ligand complex using the ProLIF package^48^, specifically evaluating hydrogen bonds, halogen bonds, salt bridges, and π-π tacking interactions. Anchors were defined as pocket atoms within a 4 Å distance cutoff from any ligand atom.

In the post-training stage, each ligand was also converted into its DSMILES representation and then was inputted into the decoder. Crucially, the encoder received binding pocket representations as the input at this stage. To enable multi-scenario molecular generation capabilities, training sequences underwent algorithmic enhancement using both stochastic (Supplementary Algorithm S3) and fragment-retained (Supplementary Algorithm S4) augmentation algorithms. UniLingo3DMol also maintained teacher-forced autoregressive reconstruction method for producing target ligands to specified binding pockets.

#### 4.3.4. Fine-tuning Stage

While the post-training stage enabled ligand generation, optimal performance for UniLingo3DMol necessitated a subsequent fine-tuning stage. This involved loading the post-training parameters and retraining the model on high-quality experimental pocket-ligand complex structures (RComplex database Level 0 data). The DUD-E protein targets served as a homology filter, preventing information leakage. Critically, pocket definitions, NCIs, and anchors were derived using the identical methodology established in the post-training phase, and the same training strategy was employed.

#### 4.3.5. *De Novo* Data and Fragment-retained Data used in UniLingo3DMol Training

The multi-stage training strategy of UniLingo3DMol relied on distinct types of molecular data, categorized as *de novo* data and fragment-retained data, to equip the model for different molecular design scenarios. The composition and application of these data types varied across the pre-training, post-training, and fine-tuning stages.

In the pre-training stage, the training data consisted exclusively of *de novo* data, meaning complete molecules without any pre-specified fragments to be retained. This stage focused on building a foundational understanding of chemical language and 3D structure from a vast dataset of commercially accessible compounds.

For the post-training and fine-tuning stages, the training data was algorithmically augmented to include both *de novo* and fragment-retained data. This was critical for achieving multi-scenario capability. The stochastic augmentation algorithm (Supplementary Algorithm S3) generated data suitable for *de novo* and single-fragment-retained design, while the specialized fragment-retained augmentation algorithm (Supplementary Algorithm S4) created data mimicking the more complex connectivity patterns of multiple-fragment-retained design. Consequently, during these stages, UniLingo3DMol learned to generate molecules conditioned on a binding pocket, both from scratch (*de novo*) and while satisfying specific constraints to retain one or more molecular fragments.

#### 4.3.6. Various Training Tasks

To address the challenge of designing molecules using a single model in various scenarios with different inputs, we introduced a multi-task training strategy to train UniLingo3DMol. Specifically, we defined three training tasks during the post-training and fine-tuning stages:

- **Task1**: NCI and anchor prediction based on given binding pocket structures.
- **Task2**: unconstrained molecular generation without pocket or NCI and anchor specifications.
- **Task3**: conditional molecular generation guided by specified binding pockets with NCI and anchor constraints.

These tasks require different inputs, which also require UniLingo3DMol to adaptively handle various inputs. To achieve this goal, UniLingo3DMol underwent special designs detailed in the following Methods Section 4.4. To maximize the effectiveness of each task, the above three tasks were iteratively trained at each epoch during training. The pseudocode of the above training process was shown in Supplementary Algorithm S2.

### 4.4. Model Architecture Across Multiple Scenarios

UniLingo3DMol was built on the transformer-based structure under the encoder-decoder architecture. Below, we introduced how UniLingo3DMol adapted to handle different inputs and generate outputs in various scenarios.

#### 4.4.1. Encoder

During the pre-training stage, the encoder processed a noise-perturbed 3D molecular structure comprising element types and spatial coordinates. Formally, the perturbed molecule was represented as 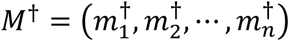, where each heavy atom contains both chemical and geometric information: 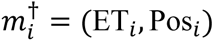 with Pos_*i*_ = (*x*_*i*_, *y*_*i*_, *z*_*i*_) ∈ *R*^3^. The position feature 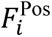 was generated through a multi-layer perceptron (MLP) that processed Euclidean coordinate embeddings, and then the input feature 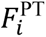 was formally represented as:

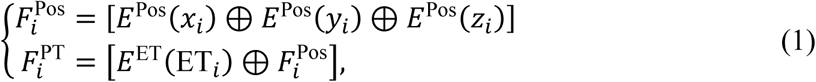

where

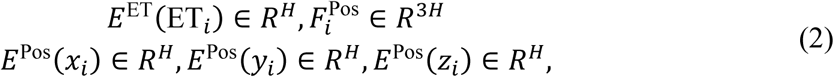

where *E*^ET^ and *E*^Pos^ denote the learnable embedding functions for the element type, and the corresponding coordinates, respectively. *H* is the size of the embedded vectors. The symbol “+” represents the element-wise addition operator, and the symbol “ [⊕] “ represents the concatenation operator.

During the post-training and fine-tuning stages, the input modality transitioned from perturbed molecular structures to protein binding pockets. To address this domain adaptation challenge and enable dynamic feature alignment across input scenarios, we introduced an MLP projection layer for feature space alignment. Specifically, the input feature representation *F*^PoT-FT^ was explicitly designed to handle multi-task scenarios through task-dependent feature composition. For instance, *F*^PoT-FT^ in Task3 was formulated as:

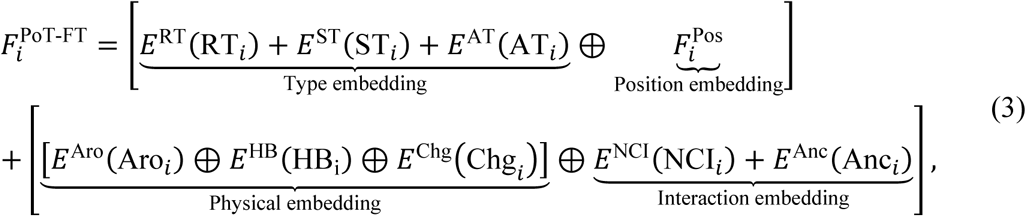

Due to the unavailability of NCI data and anchor information, we revised the input feature representations for Task1 and Task2 as follows:

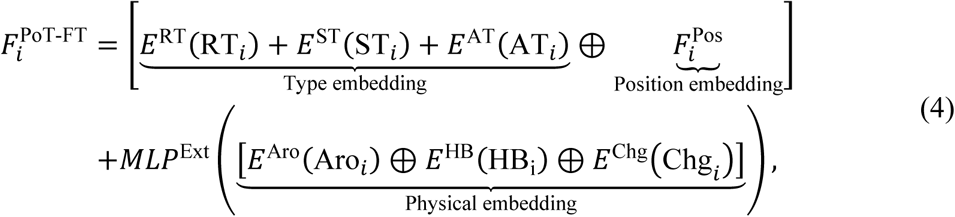

where

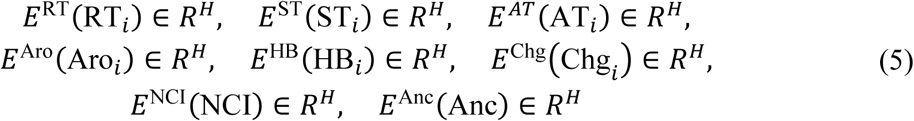

Here, the learnable embedding functions *E*^RT^ (residue type), *E*^ST^ (symbol type), *E*^*AT*^ (atomic type), *E*^Aro^ (aromaticity), *E*^HB^ (hydrogen bond donor and acceptor), *E*^Chg^ (formal charge), *E*^NCI^ (NCI sites) and *E*^Anc^ (anchor sites) collectively process the pocket’s features, while an MLP performs the nonlinear transformation from *R*^3*H*^ to *R*^4*H*^ through hidden layer computations.

Both feature representations, *F*^PT^ and *F*^PoT-FT^, first underwent sinusoidal positional encoding (PE) to capture sequential relationships, then were processed by a 6-layer transformer-based encoder with multi-head attention mechanisms, where residual connections and layer normalization were applied at each layer. Through this hierarchical processing, the pocket encoder ultimately generated the refined latent space representation *H*^Enc^, which captured both structural and contextual information of the input.

#### 4.4.2. Pocket Prediction Head

Although UniLingo3DMol demonstrated the ability to perform molecular design without giving NCI and anchor information (Task2), we proposed that integrating a dedicated NCI and anchor prediction head based on the latent space representation *H*^Enc^ could improve its capability. This auxiliary head consists of two independent MLP projections, one used to enhance pocket feature representation by identifying key binding sites in the pocket, and the other used to identify potential anchor sites to establish spatial constraints and inform the ligand generation process through clear interaction perception guidance. The above process is described as:

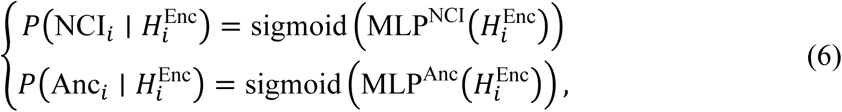

where sigmoid denotes the sigmoid function. Since the outputs of 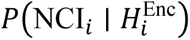 and 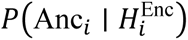 are probabilities in the range [0, 1], we binarized these probabilities using a threshold of 0.5 to determine whether each heavy atom in the pocket is an NCI or anchor site.

#### 4.4.3. Decoder

The ligand decoder adapted the same architecture as the pocket encoder, and the input representation was formally defined through a composite embedding scheme:

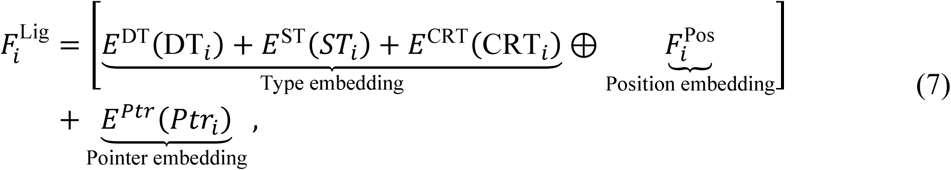

where the embedding dimensions satisfy:

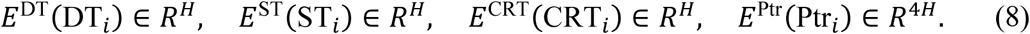

Here, the learnable embedding operators {*E*^DT^, *E*^ST^, *E*^CRT^, *E*^Ptr^} and 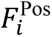 generated distributed representations capturing molecular atom-level 2D topological connectivity in chemical graphs and molecular geometry with 3D spatial conformations.

Due to the use of autoregressive training schema in UniLingo3DMol, the ligand feature *F*^Lig^ was initially transformed via sinusoidal positional embedding, then propagated through a 12-layer transformer-based decoder. This decoder integrated dual attention pathways: (1) self-masked attention for autoregressive context modeling of intra-ligand feature dependencies, and (2) cross-modal attention for iteratively aligning ligand semantics with the pocket encoded latent space *H*^Enc^. The processed hierarchy culminated in the final decoded latent space representation *H*^Dec^.

#### 4.4.4. Ligand Generation Heads

Building upon the latent space representation *H*^Dec^, we designed three specialized decoding heads for molecular generation: ligand token head, pointer head, and position head. The goal of the ligand token head was to predict the DSMILES type at step *i* based on last step latent space 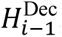, using an MLP with vocabulary projection:

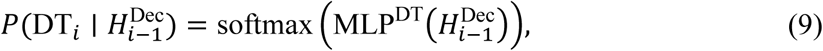

where DT_*i*_ ∈ *V*_DT_ represents the DSMILES token type at each decoding step *i*. During training, we calculated its embedding *E*^DT^(DT*_i_*) in a teacher-forced manner. At inference, we instead sample DT*_i_* via multinomial sampling to enhance molecular diversity, subsequently generating its corresponding embedding through the learned embedding function *E*^DT^.

Although DT*_i_* was obtained via the ligand token head, its connectivity with other previous tokens was still unclear. Therefore, the ligand pointer header was used to predict positional connections for a given DSMILES token. We designed this head such that connectivity prediction is conditioned on the known DSMILES type, based on the following formulation:

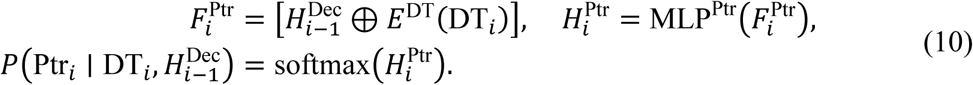

Once the DSMILES token type and its positional pointer were known, its topology was determined. To predict its spatial coordinates, the ligand position head was used together with four MLP projections. Here, we established two coordinate systems to predict its 3D coordinates: the global coordinate system, using Euclidean coordinates (*x*, *y*, *z*); the local coordinate system, using relative spatial positions such as bond lengths, bond angles, and dihedral angles (*r*, *θ*, *ϕ*). Consistent with the ligand pointer head, coordinate prediction was conditioned on the known topological structure.

We adopted a local-to-global inference order: local coordinates were predicted first, followed by global coordinates. This design was motivated by the fact that the local coordinate system involves fewer degrees of freedom than the global one. By progressing from lower to higher complexity, the model refines its coordinate estimates in a stepwise manner. The overall formulation is described as follows:

The local coordinate system:

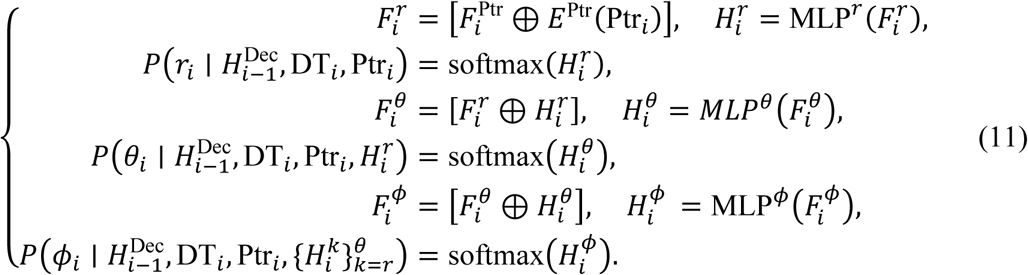

The global coordinate system:

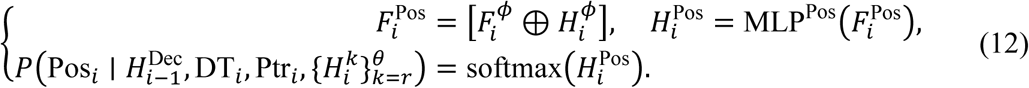

### 4.5. Multi-Task Driven Molecular Generation in UniLingo3DMol

The multi-task training architecture of UniLingo3DMol enabled molecule generation through distinct task configurations under varying input conditions, specifically with or without NCI and anchor site specifications.

#### Generation with given NCI and anchor sites

When provided with explicit NCI and anchor sites, UniLingo3DMol leveraged Task3 to navigate the chemical design space. This approach improved the proportion of generated molecules that satisfy NCI criteria within specific protein binding pockets.

#### Generation without NCI and anchor sites

For the scenario lacking prior NCI and anchor information, two generation pathways were available. For the first option, we directly employed Task2 to generate molecules without prior information. For the second option, we built a cascaded workflow employing Task1 (NCI and anchor prediction) followed by Task3. The comparison of these two pathways is shown in Supplementary Table S5, and experimental results suggested that the first option achieved better performance than the second performance. We believed that this performance came from end-to-end training for Task2.

#### 4.5.1. DSMILES Token Prediction

The ligand generation followed an autoregressive sequence where DSMILES tokens were progressively determined through discrete probabilistic selection. At each decoding step *i*, UniLingo3DMol computed token-wise probability distributions over the DSMILES vocabulary through the ligand token head. These probabilistic outputs drove a multinomial sampling operation to determine the DSMILES token type at step *i*.

#### 4.5.2. Token’s Pointer Prediction

At each decoding step *i*, after identifying the DSMILES token type, we proceeded to predict its topological connectivity by using the ligand pointer head. Based on the positional pointer definitions introduced in Methods Section 4.2, invalid pointer positions were systematically excluded by applying connectivity constraints. Specifically, non-asterisk tokens were restricted to point to their own positions (self-reference), while asterisks tokens were permitted to connect only to preceding asterisk tokens within the sequence. The full strategy is for positional pointer prediction formally described in Supplementary Algorithm S5, which outlines the constraint-based selection process for valid pointer assignments.

#### 4.5.3. Token’s Coordinates Prediction

Here, we predicted the coordinates of the DSMILES token at each step *i* using the ligand position head. Leveraging the established local and global coordinate systems, the local coordinates (*r*, *θ*, *ϕ*) was used to help the prediction of global coordinates (*x*, *y*, *z*). Similar to the approach in Lingo3DMol^8^, the coordinates of the token were determined in the search space that was defined by its local coordinates. This search space is defined as follows:

- *r* ± 0.1 Å (angstroms) for bond lengths.
- *θ* ± 2 ^∘^ (degrees) for bond angles.
- *ϕ* ± 2 ^∘^ (degrees) for dihedral angles.

Within the defined search space, we identified the coordinates corresponding to the maximum joint probability from the predicted global coordinate distributions.

The above processes were cyclically performed until the generation was finalized, with the corresponding pseudocode provided in Supplementary Algorithm S6. In addition, a schematic overview of the two-fragment-retained molecular generation workflow is illustrated in Extended Data Figure 7.

### 4.6. RComplex Database Construction

The RComplex database was constructed to support UniLingo3DMol training through a multi-stage computational pipeline designed for the extraction and modeling of active compound-protein complex structures (Extended Data Figure 1).

The initial phase involved data acquisition and structural curation. A total of 48,222 protein–ligand complexes were retrieved from the Protein Data Bank^33^ using a multi-step filtering process. First, crystal structures containing multiple proteins associated with different UniProt IDs were excluded, retaining only single protein–ligand complexes. Heteromolecules were then classified as valid ligands or non-relevant compounds based on predefined PDBbind^49^ lists and frequency analysis; molecules present in over 100 crystal structures were eliminated to remove common solvents, lipids, and frequently appearing co-factors. Additionally, metal-containing molecules were systematically discarded. Further manual curation excluded known cofactors and impurities, ensuring only genuine ligands of interest were included.

Subsequently, bioactive ligand data were curated from ChEMBL^34^, PubChem^35^, and GOSTAR^36,37^. Only compounds possessing target information (UniProt ID) and reported bioactivity (K*_d_*, IC_50_, K*_i_*, or EC_50_) no worse than 10 µM were utilized. These ligands, initially associated with one or more UniProt IDs, were then matched to specific PDB IDs. For all compounds associated with a given PDB ID, molecular docking was performed against the protein pocket extracted from that PDB ID using Glide in Standard Precision (SP) mode. Constraints were applied based on the maximum common substructure shared with the native co-crystallized ligand during docking. Each docking campaign generated up to 5 ligand conformations within the respective binding site. The docking-generated complex structures subsequently underwent a stringent computational filtering process. Complex structures were retained only if they satisfied the following criteria: a Glide SP docking score ≤ −10 kcal/mol and an MM/GBSA binding free energy ≤ −50 kcal/mol (calculated using the Prime MM-GBSA method^50,51^).

All the protein-ligand complex structures, encompassing both curated structures from PDB and filtered docking structures, were then classified into three distinct confidence levels. Level 0 designated the previously curated 48,222 native protein-ligand crystal structures. For all filtered docking structures, each associated with a unique PDB ID, they were classified based on their structural relationship to the native co-crystallized ligand for that PDB ID. Level 1 included compounds exhibiting a reciprocal substructure relationship (where either the bioactive molecule or the co-crystallized ligand is a substructure of the other) and a Tanimoto similarity ≥ 0.5. Level 2 comprised compounds meeting only a Tanimoto similarity threshold of ≥ 0.5. This entire integration and classification process collectively yielded a consolidated dataset of 3.7 million protein-ligand complex structures, forming the RComplex database.

### 4.7. In-vitro and In-vivo Assays

#### 4.7.1. Expression and Purification of Human CBL-B N-domain

The gene encoding human CBL-B (UniProt code: Q13191) N-domain (Gln38-Asp427) was subcloned into a pET-28a vector fused with an 8xHis tag followed by an HRV 3C protease site at the N-terminus. The plasmids were transformed into E. coli SHuffle T7 competent cells (New England Biolabs). The cells were cultured in Luria Broth (LB) medium in a 37 °C incubator shaking at 220 rpm. When OD600 reached 0.8, the medium was supplied with 0.5 mM isopropyl β-D-thiogalactoside (Sigma). After culturing for another 22 hours at 16 °C, the cells were collected by centrifugation and were stored at - 80 °C.

The cell pellets were resuspended in buffer A (20 mM Tris pH 8.0, 100 mM NaCl, 2 mM β-mercaptoethanol, and 10% glycerol) supplemented with protease inhibitors. After cell breaking by ultrasonication, the cell lysate was centrifugated for 30 min at 45000 rpm. Subsequently, the supernatant was transferred and loaded onto Ni-NTA gels in a gravity column. The resin beads were washed by 10 column volume (CV) of buffer A supplemented with 35 mM imidazole. The protein samples were eluted by 15 mL elution buffer (20 mM Tris pH 8.0, 100 mM NaCl, 2 mM β-mercaptoethanol, and 300 mM imidazole). The His tag was removed by PreScission protease (1:50 w/w ratio) digestion for 2 h on ice. The samples were concentrated and loaded onto a Superdex 200 Increase 10/300 GL column (Cytiva) preequilibrated in buffer containing 20 mM Tris, 150 mM NaCl, 5 mM β-mercaptoethanol, pH 8.0. The peak fractions were pooled and concentrated to 20-22 mg/mL. Aliquots of the concentrated samples were flash frozen in liquid nitrogen and stored at -80 °C.

#### 4.7.2. Crystallization, Data Collection and Structure Determination

Before crystal screening, the CBL-B N-domain samples were supplemented with 1 mM Cmpd. 7 and incubated on ice for 1 h. Initial crystals were observed in CrystalScreen Kit G6 (0.1 M HEPES pH 7.5, 10% w/v Polyethylene glycol 6,000, 5% v/v (+/-)-2-Methyl-2,4-pentanediol). This crystallization condition was optimized. Finally, 1μL protein solution was mixed with 1μL reservoir solution (0.1 M HEPES pH 7.3, 7% w/v polyethylene glycol 6,000, 5% v/v (+/-)-2-Methyl-2,4-pentanediol) for each drop, and large crystals were obtained within 7 days at 18 °C. Crystals were transferred into cryoprotectant containing mother reservoir solution supplemented with 25% ethylene glycol, then flash frozen in liquid nitrogen.

X-ray diffraction data of Cmpd. 7-CBL-B crystals were collected at beamline BL19U1 of the Shanghai Synchrotron Radiation Facility (SSRF). X-ray diffraction data were integrated and scaled using the program HKL-2000^52^. The initial phase was obtained by molecular replacement (MR) with the CBL-B structure (PDB code: 8GCY) as a reference model using the program Phaser in PHENIX^53^. Several rounds of manual model correction and refinements were performed iteratively using COOT and PHENIX refinement. Statistics for data collection and processing are shown in Extended Data Table 1.

#### 4.7.3. CBL-B TR-FRET Assay

The biochemical activity of compounds against human CBL-B was determined using a time-resolved fluorescence resonance energy transfer (TR-FRET) assay, which utilized 30 nM SRC substrate. First, 10 µL of each test compound were transferred to a 384-well low dead volume (LDV) Echo plate. Second, compounds were diluted and transferred 150 nL to the assay plate using an Echo liquid handler, establishing 10 concentration points in duplicate. A 3.5 nM solution of human CBL-B enzyme, prepared in assay buffer, was then added at 5 µL per well to the assay plate; this was followed by centrifugation at 1500 rpm for 1 minute and a 1-minute preincubation at 25 °C. The reaction was initiated by adding 5 µL of a prepared substrate mixture to each well of the Corning 784075 assay plate, after which the plate was centrifuged again at 1500 rpm for 1 minute and incubated for 2 hours at 25 °C. Following this incubation, 5 µL of a binding assay mix was added to each well, and the plate underwent a final centrifugation at 1500 rpm for 1 minute before being incubated for an additional 60 minutes at 25 °C. Finally, the TR-FRET signal at 520/620 nm was recorded using an Envision plate reader, and the 50% inhibitory concentration (IC_50_) values were calculated from the recorded data using the log (inhibitor) vs. normalized response–variable slope model provided by GraphPad Prism software.

#### 4.7.4. Jurkat IL-2 Induction Assay

Jurkat E6-1 cells were cultured in RPMI 1640 medium supplemented with 10% fetal bovine serum (FBS). For the assay, 100 μL of cell suspension (containing 1×10^5^ cells) were seeded into each well of 96-well plates. The cells were then treated with varying concentrations of compounds, in triplicate, for 60 minutes. Following compound treatment, the cells were stimulated for 48 hours with 0.5 μg/mL anti-human CD3 antibody and 0.5 μg/mL anti-human CD28 antibody. Subsequently, cell culture supernatants were harvested, and the concentration of interleukin-2 (IL-2) was quantified using an enzyme-linked immunosorbent assay (ELISA) kit.

#### 4.7.5. CT26 Murine Syngeneic Model Efficacy Studies

All animal studies were conducted at ProOnco Therapeutics Co., Ltd., under protocols approved by and in compliance with the Institutional Animal Care and Use Committee (IACUC) of the institution. CT26 tumor cells were cultured in vitro using RPMI 1640 medium supplemented with 10% fetal bovine serum (FBS) at 37 °C with 5% CO_2_. For tumor induction, BALB/c mice were subcutaneously inoculated in the right flank with 3×10^5^ CT26 tumor cells suspended in 0.1 mL of phosphate-buffered saline (PBS). Drug treatments were initiated when tumor volumes reached between 50-70 mm^3^. All enrolled tumor-bearing mice were randomized into study groups based on both tumor size and body weight. The investigational compound, Cmpd. 20, was administered orally as a homogeneous suspension once daily (QD), while anti-mPD-1 antibody (Clone RMP1-14, Bio X Cell) was administered intraperitoneally (i.p.) as a solution twice a week (BIW). Study animals were monitored daily for signs of morbidity and mortality. Body weight and tumor volumes were measured every 3 days. Tumor volumes were determined in two dimensions using a caliper and calculated in mm^3^ using the formula: V = (L ×W ×W) / 2, where V represents tumor volume, L is the longest tumor dimension (length), and W is the longest tumor dimension perpendicular to L (width). Individual animals were euthanized when their tumor volume exceeded 2,000 mm^3^, and the entire experiment was terminated when the mean tumor volume of the control group surpassed this threshold.

## Data and Code Availability

The DUD-E dataset is publicly available at https://dude.docking.org/. Our source code is deposited in Code Ocean. The generated molecules, including their evaluation metrics, and a subset of the training data are deposited in Figshare. All such deposited resources are available for peer review and will be made publicly accessible upon manuscript publication.

## Acknowledgements

This work was supported by the Beijing Municipal Science and Technology Commission (grant no. Z241100007724005 received by B.H.). This study was also funded by the National Key R&D Program of China (grant no. 2022YFF1203004 received by B.H.).

## Author Contributions

B.H. and H.W. conceived the study. H.W. build the AI model. G.S, M.Y., and Y.G. synthesize and evaluated the compounds. W.Z. provided instructions for artificial intelligence modelling. F.F. and B.Z prepared the training data. H.W. provided instructions for compound evaluation. B.H. and Z.L. provided instructions on evaluation framework. D.J. provided instructions for structural biology study. Y.Z., W.F., Y.X., B.X., Y.W., X.X., Y.W., and C.L. supported the model evaluation.

