## Extended Data Figures and Tables for "A unified language model bridging *de novo* and fragment-based 3D molecule design delivers potent CBL-B inhibitors for cancer treatment"

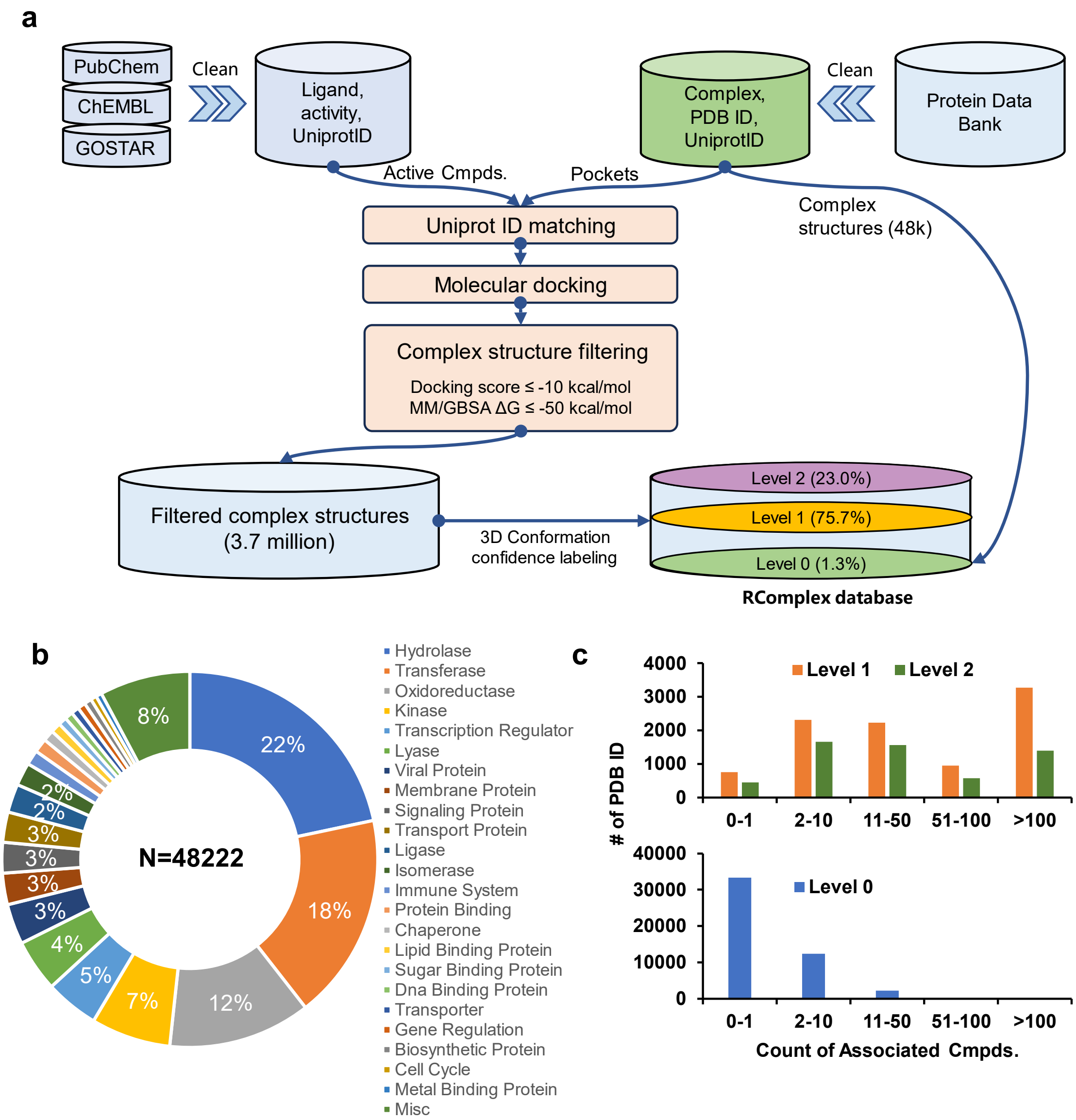

**Extended Data Figure 1. RComplex database construction.**

**(a)** Database construction procedure. The RComplex database contains 3.7 million complex structures, each associated with a unique PDB ID indicating the source of the pocket structure in that complex. These complexes are classified into three distinct confidence levels based on ligand similarity to the co-crystallized ligand of that PDB ID: Level 0 for identical ligands, Level 1 for reciprocal substructure relationships with a Tanimoto similarity ≥ 0.5, and Level 2 for compounds meeting only a Tanimoto similarity threshold of ≥ 0.5. These confidence levels account for 1.3%, 75.7%, and 23% of the RComplex database volume, respectively. Details are provided in Methods Section 4.6.

**(b-c)** RComplex database profile. A total of 48,222 PDB structures were curated to form the pocket basis for the RComplex database. Panels **(b)** and **(c)** illustrate the distribution of these PDB IDs by target type and their associated ligand count within RComplex, respectively.

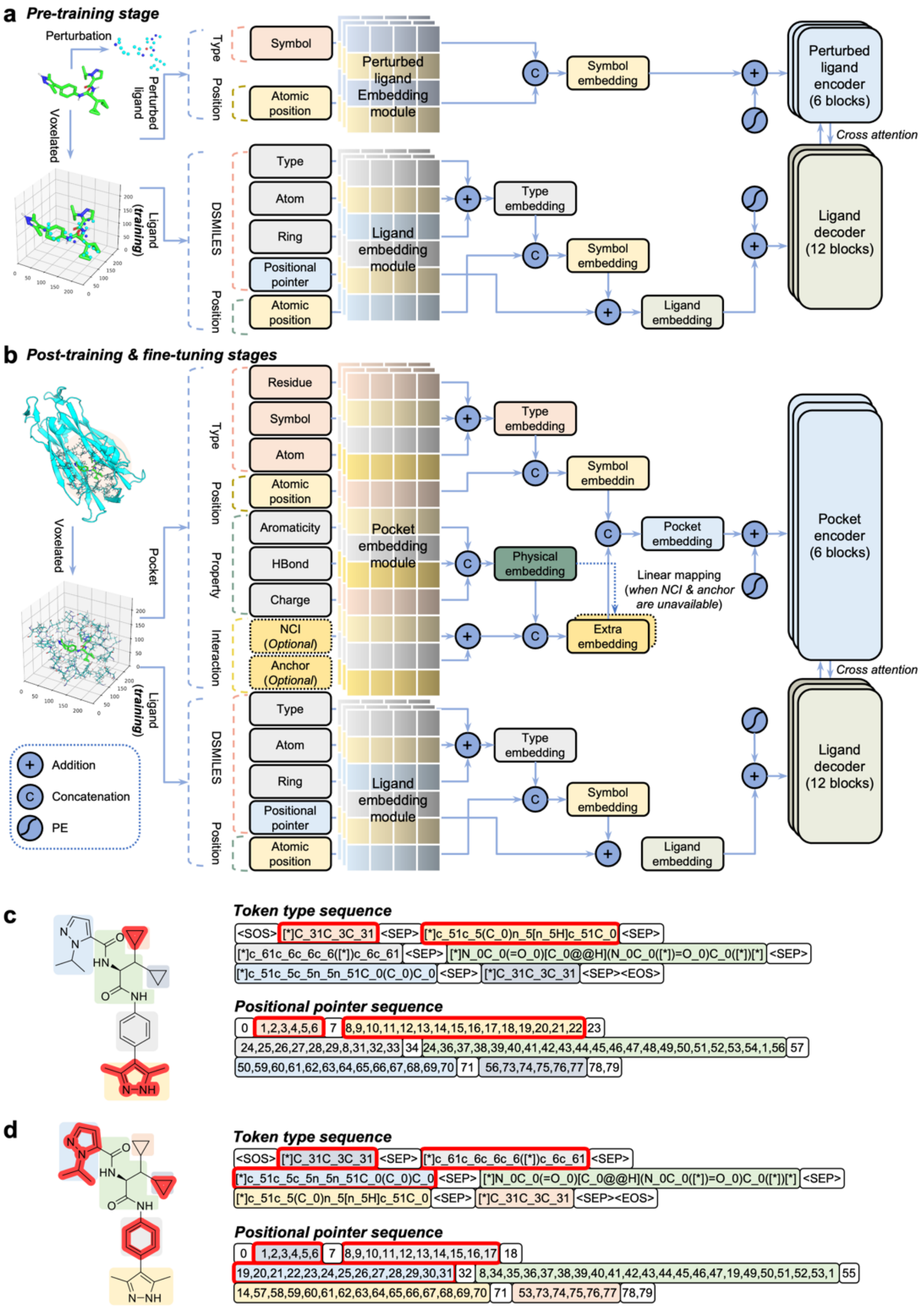

**Extended Data Figure 2. Working mechanism of UniLingo3DMol across training stages.**

**(a) Pre-training stage.** The encoder processes perturbed ligand inputs, while the decoder generates ligands as DSMILES sequences with restored 3D conformations.

**(b) Post-training and fine-tuning stages.** Here, the encoder receives protein binding pockets as input.

**(c-d) DSMILES representations of molecules with retained fragments.** Molecules with two **(c)** or three **(d)** unconnected fragments retained (highlighted in red) can be expressed using DSMILES. In these representations, the retained fragments are positioned at the beginning of the sequence, which allows for the subsequent autoregressive generation of additional atoms.

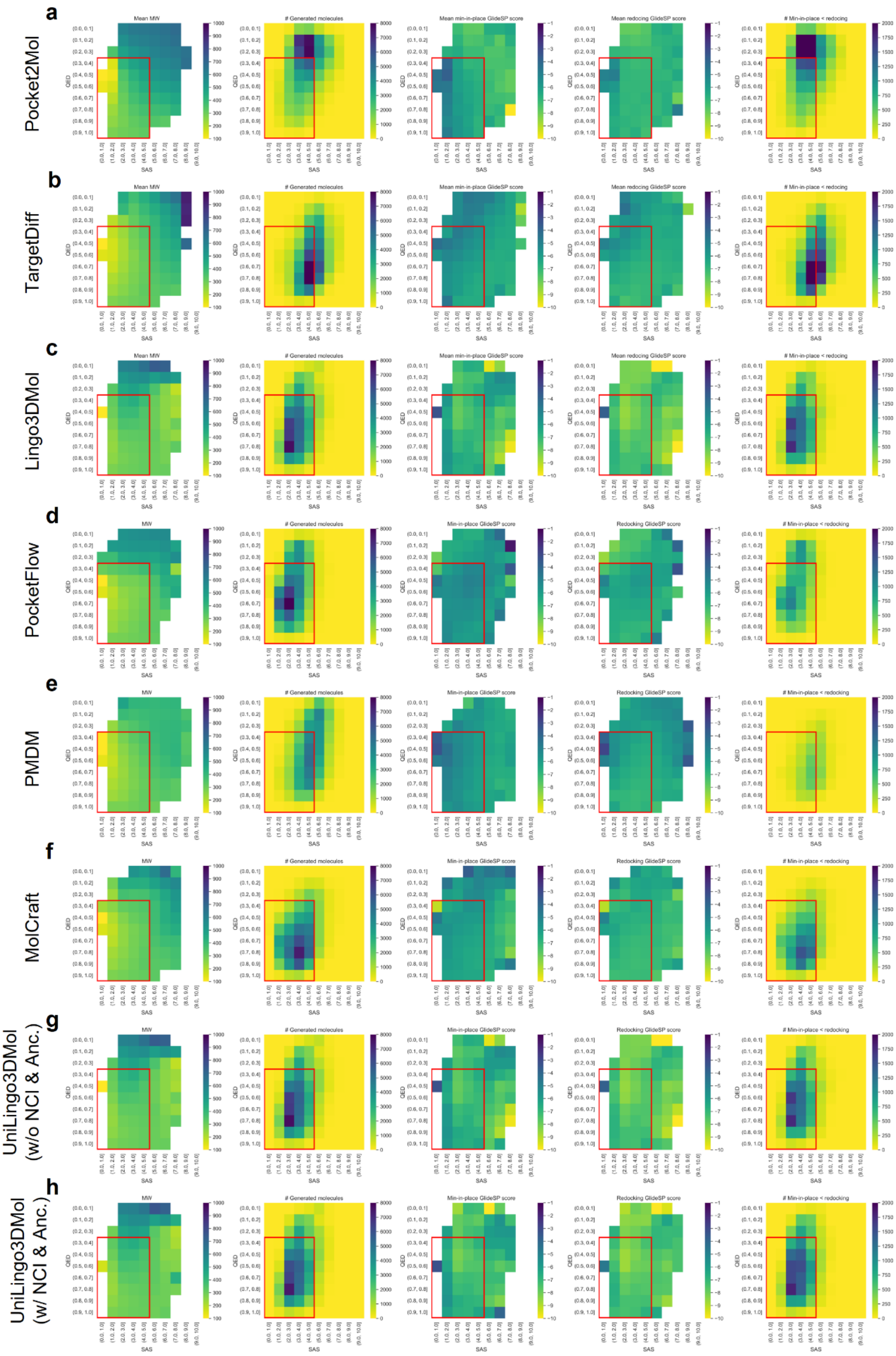
a
Pocket2Mol
b
TargetDiff
c
Lingo3DMol
d
PocketFlow
e
PMDM
f
MolCraft
g
UniLingo3DMol (w/o NCI & Anc.)
h
UniLingo3DMol (w/ NCI & Anc.)
Mean MW
MW
### Generated molecules
Mean min-in-place GlideSP score
Mean redocking GlideSP score
Min-in-place GlideSP score
Redocking GlideSP score
### Min-in-place < redocking
QED
SAS

**Extended Data Figure 3. Distributions of molecules generated by various models on the *de novo* design evaluation set.**

The drug-like region with QED > 0.3 and SAS < 5 is indicated with red boxes. (**a-h)** show the distributions on molecules generated by Pocket2Mol **(a)**, TargetDiff **(b)**, Lingo3DMol **(c)**, PocketFlow **(d)**, PMDM **(e)**, MolCraft **(f)**, UniLingo3DMol without NCI and anchor constraints **(g)**, and UniLingo3DMol with NCI and anchor constraints **(h)**, respectively. Heatmaps visualize the distribution of key properties across the generated molecules, including: (from left to right) molecular weight, counts, min-in-place GlideSP scores, redocking GlideSP scores, and the count of instances where the min-in-place GlideSP score is lower than the redocking GlideSP score. These distributions are depicted along SAS and QED.

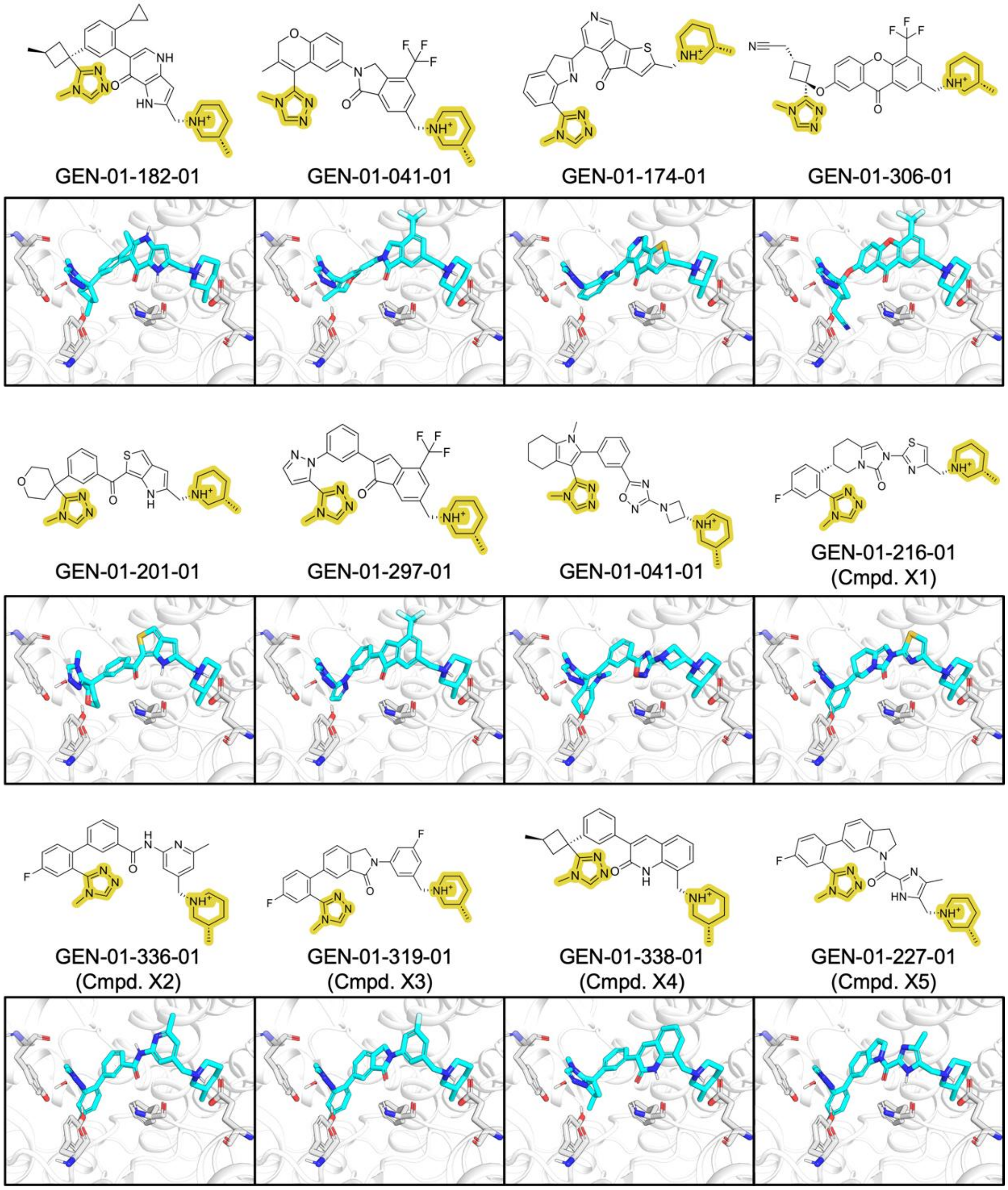

**Extended Data Figure 4. Examples of UniLingo3DMol-generated molecules demonstrating classical and novel moieties in Region B.**

Retained fragments are highlighted in yellow. Molecules GEN-01-182-01 and GEN-01-041-01 represent classical moieties in Region B (region definitions provided in Figure 3a), specifically pyrrolopyridine and trifluoromethyl-isoindolin derivatives. Other molecules show Region B moieties with varying degrees of novelty. Among them, Molecules GEN-01-174-01, GEN-01-306-01, GEN-01-201-01, and GEN-01-297-01 were eliminated due to the presence of potentially toxic structures (e.g., three-membered fused ring systems or specific ketones). Molecule GEN-01-041-01 was relatively challenging to synthesize. Cmpd. X1-X5 were selected, and to minimize synthetic complexity and cost, only their A, B, and D regions were synthesized, yielding Cmpd. 1-5 (Figure 3c).

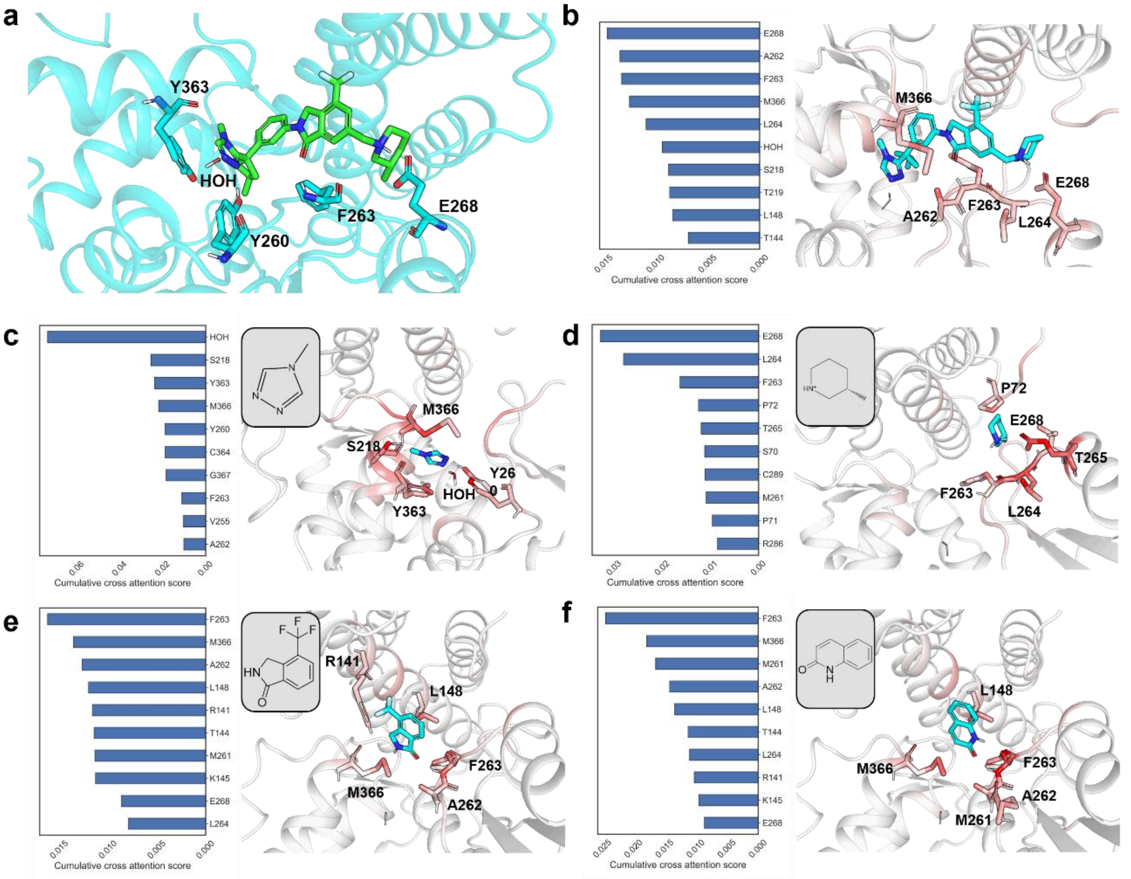

**Extended Data Figure 5. Cross-attention mechanism visualization of the UniLingo3DMol model's understanding of protein-ligand interactions.**

**(a)** Co-crystallized ligand within the CBL-B binding pocket (PDB ID: 8GCY). This complex structure serves as the context for attention mechanism analysis.

**(b-f)** Ligand and fragment-specific attention patterns within the CBL-B binding pocket. Each panel **(b-f)** displays a bar chart and a corresponding 3D visualization. Bar charts show the top 10 residue-level cumulative cross-attention scores from the entire ligand in (**b**) and specific fragments in (**c-f**). These scores aggregate the model's eight parallel attention heads, first by summing atom-level attention scores for the relevant ligand **(b)** or fragment **(c-f)** atoms, then averaging for all atoms within the same protein residue. In 3D ligand-protein complex structures, protein atoms are colored by their individual atomic attention scores, with a gradient ranging from white (low attention) to red (high attention). The top 5 residues from each corresponding bar chart are explicitly labeled and depicted as sticks in the 3D visualization.

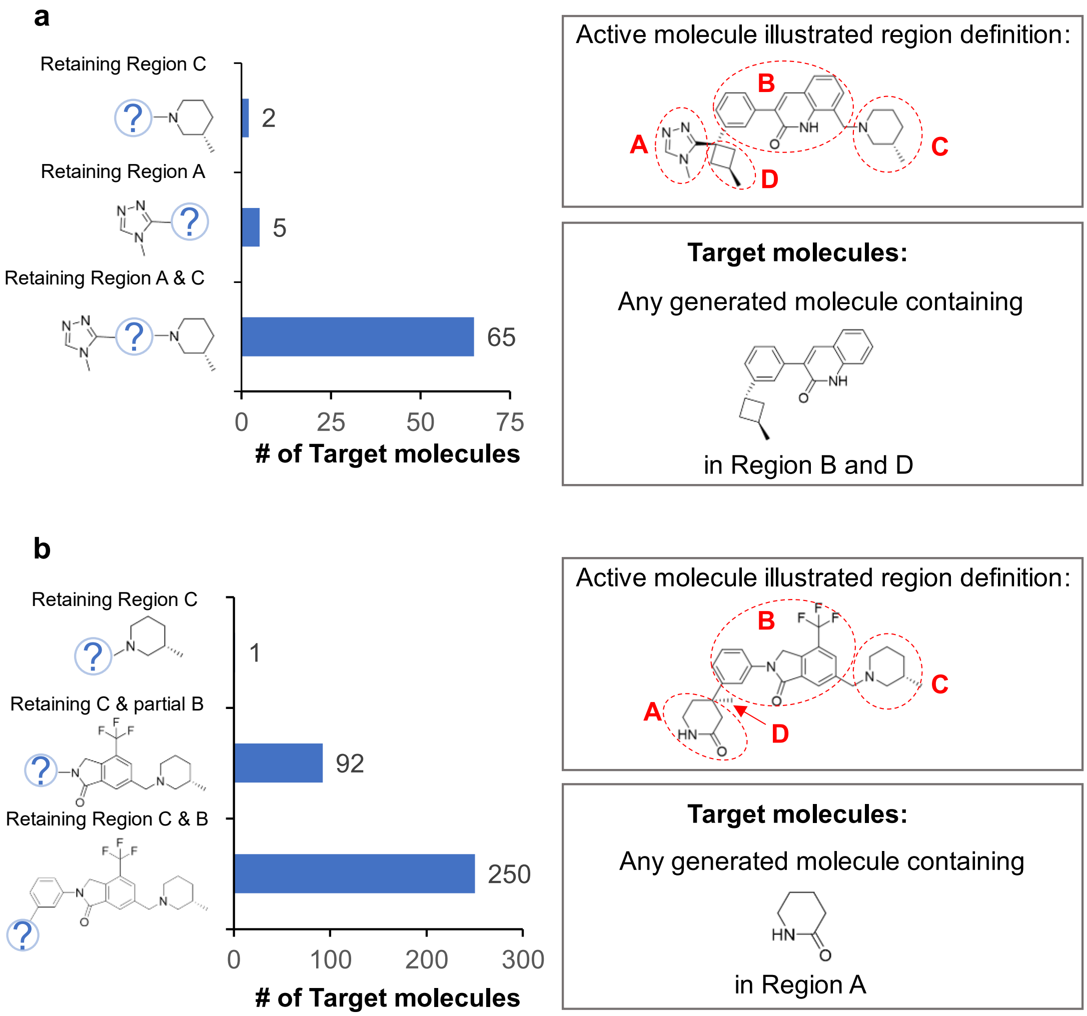

**Extended Data Figure 6. Impact of retained fragments on the generation of specific target molecules.**

Each panel displayed the count of target molecules generated by UniLingo3DMol, sampled up to 3 million times with varying retained fragments. The structure of a relevant active compound is provided to illustrate region definition.

**(a)** Impact of varying retained fragments on generating molecules with target scaffold in Region B and D.

**(b)** Impact of varying retained fragments on generating molecules with specific moieties in Region A.

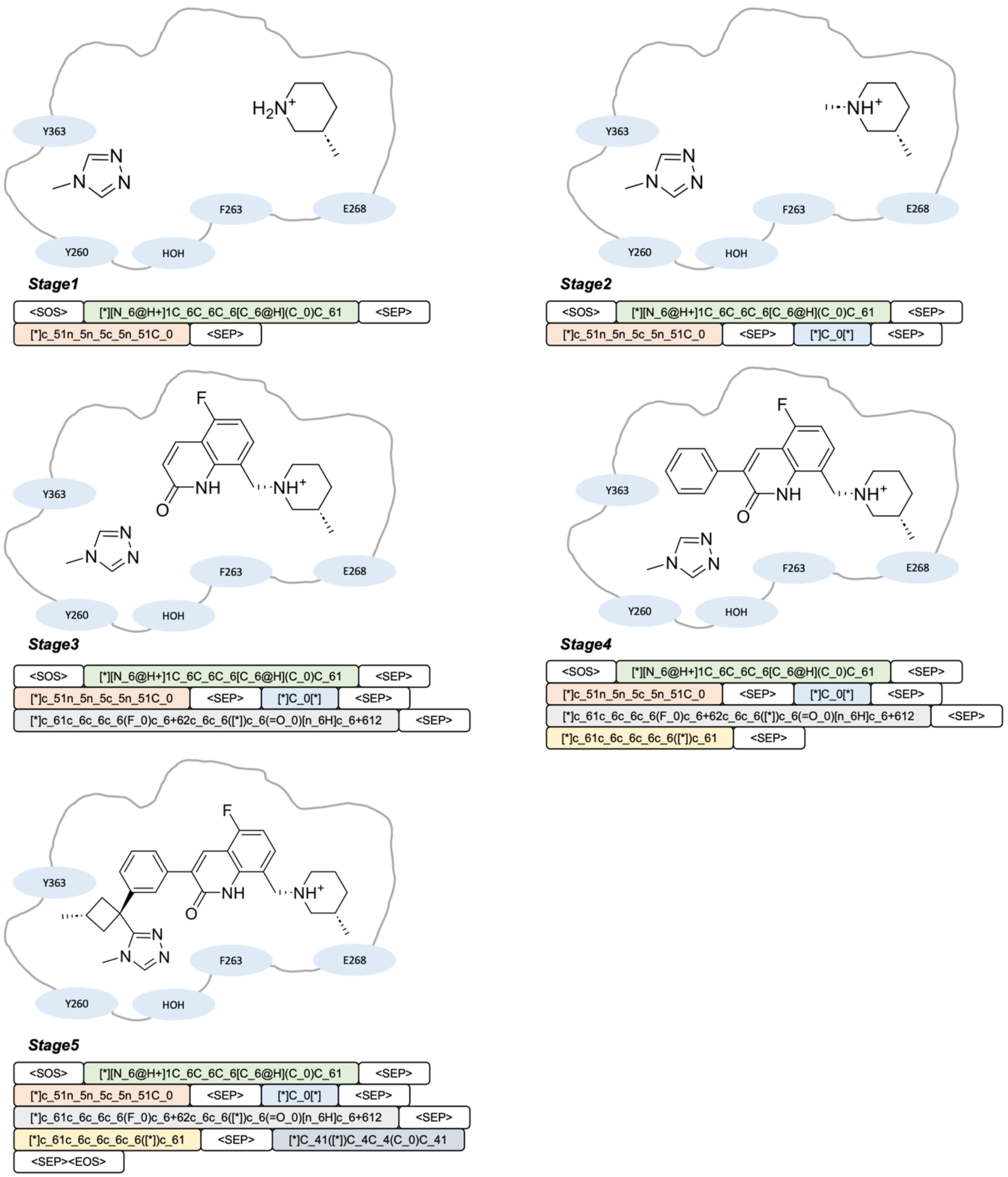

**Extended Data Figure 7. Visualization of molecular generation stages with UniLingo3DMol in the two-fragment-retained scenario.**

**Extended Data Table 1. Data collection, refinement and validation statistics.**

| | Cmpd. 7-CBL-B (PDB: 9WDH) |
|---|---|
| **Data collection** | |
| **Space group** | P21 |
| **Unit Cell** | a=57.1 Å, b=102.0 Å, c=74.1 Å<br>α=γ= 90°, β=89.86° |
| **Resolution** | 2.8 Å (2.95 - 2.80) |
| **Total measured reflections** | 20,806 |
| **Completeness (%)** | 99.1 (98.0) |
| **Redundancy** | 3.2 (3.1) |
| **I/σ** | 7.2 (2.1) |
| **$R_{merge}$** | 0.172 (0.346) |
| **CC(1/2)** | 0.988 (0.964) |
| **Resolution range** | 59.9 – 2.8 Å |
| **Refinement** | |
| **Initial model** | PDB code: 8GCY |
| **R, $R_{free}$** | 25.4%, 30.7% |
| **Model composition** | |
| **Non-hydrogen atoms** | 6,483 |
| **Protein atoms** | 6,224 |
| **Ligand atoms** | 144 |
| **Wilson B-factor (Å$^2$)** | 13.4 |
| **Average B factors (Å$^2$)** | |
| **Protein** | 15.9 |
| **Ligand** | 6.2 |
| **R.m.s. deviations** | |
| **Bond lengths (Å)** | 0.004 |
| **Bond angles (°)** | 0.79 |
| **Validation** | |
| **MolProbity score** | 2.87 |
| **Poor rotamers (%)** | 2.3 |
| **Ramachandran plot** | |
| **Favored (%)** | 89.7 |
| **Allowed (%)** | 8.8 |
| **Disallowed (%)** | 1.5 |

**Extended Data Table 2. Cmpd. 20 preclinical PK and safety profile.**

| Compound ID | Cmpd. 20 |
|---|---|
| CT26 CTG assay, IC50 (µM) | 17 |
| PPB% Bound (mouse) | 99.46% |
| CYP1A2, 2C9, 2C19, 2D6, 3A4/5 inhibition, (10µM) | 3.9%, 73%, 31%, 28%, 84% |
| hERG inhibition%, (10µM) | 32.8% |
| PO. mouse PK (10mg/kg): $t_{1/2}$ (h), $C_{max}$ (ng/mL), $AUC_{0\text{-}last}$ (h.ng/mL) | 1.36, 586, 591 |
| PO. mouse PK (30mg/kg): $t_{1/2}$ (h), $C_{max}$ (ng/mL), $AUC_{0\text{-}last}$ (h.ng/mL) | 2.17, 2945, 6561 |

**Extended Data Table 3. Valid vocabularies for atomic types, ring types, and DSMILES types.**

| | | | | | |
|---|---|---|---|---|---|
| **Atomic type vocabulary** | | | | | |
| <PAD> | <SOS> | <EOS> | <SEP> | C | N |
| O | S | P | F | Cl | Br |
| I | c | n | o | s | UnkElement |
| UnkAtomType | UnkSymbol | H | + | - | / |
| \ | @ | @@ | # | = | 1 |
| 2 | 3 | 4 | 5 | 6 | ( |
| ) | [ | ] | [*] | ([*]) | <MOL> |
| **Ring type vocabulary** | | | | | |
| 0 | 3 | 3+3 | 3+4 | 3+5 | 3+5+5 |
| 3+5+6 | 3+6 | 3+6+6 | 3+7 | 4 | 4+4 |
| 4+4+4 | 4+4+5 | 4+5 | 4+5+5 | 4+5+6 | 4+6 |
| 4+6+6 | 4+7 | 4+7+7 | 4+8 | 5 | 5+5 |
| 5+5+5 | 5+5+6 | 5+5+7 | 5+6 | 5+6+6 | 5+6+7 |
| 5+6+8 | 5+7 | 5+7+7 | 5+8 | 6 | 6+6 |
| 6+6+6 | 6+6+7 | 6+6+8 | 6+7 | 6+7+7 | 6+7+8 |
| 6+8 | 6+8+8 | 7 | 7+7 | 7+8 | 8 |
| 8+8 | 8+8+8 | | | | |
| **DSMILES type vocabulary** | | | | | |
| <PAD> | <SOS> | <EOS> | <SEP> | C_0 | C_3 |
| C_3+3 | C_3+4 | C_3+5 | C_3+5+5 | C_3+5+6 | C_3+6 |
| C_3+6+6 | C_3+7 | C_4 | C_4+4 | C_4+4+4 | C_4+4+5 |
| C_4+5 | C_4+5+5 | C_4+5+6 | C_4+6 | C_4+6+6 | C_4+7 |
| C_4+7+7 | C_4+8 | C_5 | C_5+5 | C_5+5+5 | C_5+5+6 |
| C_5+5+7 | C_5+6 | C_5+6+6 | C_5+6+7 | C_5+6+8 | C_5+7 |
| C_5+8 | C_6 | C_6+6 | C_6+6+6 | C_6+6+7 | C_6+7 |
| C_6+7+7 | C_6+7+8 | C_6+8 | C_6+8+8 | C_7 | C_7+7 |
| C_7+8 | C_8 | C_8+8 | C_8+8+8 | N_0 | N_3 |
| N_4 | N_4+5 | N_4+6 | N_4+7 | N_4+8 | N_5 |
| N_5+5 | N_5+5+5 | N_5+5+6 | N_5+6 | N_5+6+6 | N_5+7 |
| N_5+8 | N_6 | N_6+6 | N_6+6+6 | N_6+6+7 | N_6+7 |
| N_6+7+7 | N_6+8 | N_7 | N_7+7 | N_8 | O_0 |
| O_3 | O_4 | O_4+6 | O_5 | O_5+5 | O_5+6 |
| O_5+7 | O_6 | O_6+6 | O_6+7 | O_7 | O_7+7 |
| O_8 | S_0 | S_3 | S_4 | S_5 | S_5+5 |

| | | | | | |
|---|---|---|---|---|---|
| S_5+6 | S_6 | S_6+6 | S_7 | S_8 | P_0 |
| P_5 | P_6 | P_7 | F_0 | Cl_0 | Br_0 |
| I_0 | c_0 | c_3 | c_4 | c_4+6 | c_5 |
| c_5+5 | c_5+5+6 | c_5+6 | c_5+6+6 | c_5+6+7 | c_5+6+8 |
| c_5+7 | c_5+7+7 | c_5+8 | c_6 | c_6+6 | c_6+6+6 |
| c_6+6+7 | c_6+6+8 | c_6+8+8 | c_6+7 | c_6+7+7 | c_6+8 |
| c_7 | c_7+7 | n_0 | n_5 | n_5+5 | n_5+5+6 |
| n_5+6 | n_5+6+6 | n_5+6+7 | n_5+7 | n_5+7+7 | n_5+8 |
| n_6 | n_6+6 | n_6+6+6 | n_6+7 | n_6+7+7 | n_6+8 |
| n_7 | o_5 | o_6 | s_5 | s_6 | UnkElement |
| UnkAtomType | UnkSymbol | H | <MOL> | + | - |
| / | \ | @ | @@ | # | = |
| 1 | 2 | 3 | 4 | 5 | 6 |
| ( | ) | [ | ] | [*] | ([*]) |
